# Chromatin Accessibility and Pioneer Factor FOXA1 Shape Glucocorticoid Receptor Action in Prostate Cancer

**DOI:** 10.1101/2023.03.03.530941

**Authors:** Laura Helminen, Jasmin Huttunen, Niina Aaltonen, Einari A. Niskanen, Jorma J. Palvimo, Ville Paakinaho

## Abstract

Treatment of prostate cancer relies predominantly on the inhibition of androgen receptor (AR) signaling. Despite the initial effectiveness of the antiandrogen therapies, the cancer often develops resistance to the AR blockade. One mechanism of the resistance is glucocorticoid receptor (GR)-mediated replacement of AR function. Nevertheless, the mechanistic ways and means how the GR-mediated antiandrogen resistance occurs have remained elusive. Here, we have discovered several crucial features of GR action in prostate cancer cells through genome-wide techniques. We detected that the replacement of AR by GR in enzalutamide-exposed prostate cancer cells occurs almost exclusively at pre-accessible chromatin sites displaying FOXA1 occupancy. Counterintuitively to the classical pioneer factor model, silencing of FOXA1 potentiated the chromatin binding and transcriptional activity of GR. This was attributed to FOXA1-mediated repression of the *NR3C1* (gene encoding GR) expression *via* the corepressor TLE3. Moreover, the small-molecule inhibition of coactivator p300’s enzymatic activity efficiently restricted GR-mediated gene regulation and cell proliferation. Overall, we identified chromatin pre-accessibility and FOXA1-mediated repression as important regulators of GR action in prostate cancer, pointing out new avenues to oppose steroid receptor-mediated antiandrogen resistance.

## INTRODUCTION

Steroid receptors, such as the glucocorticoid (GR) and the androgen receptor (AR), belong to the nuclear receptor superfamily, and they function as hormone-activated transcription factors (TFs) involved in several crucial biological processes ranging from development to general homeostasis (1, 2). The binding of their cognate hormones promotes nuclear translocation enabling the receptor to engage with chromatin, ultimately leading to the modulation of gene expression. The modulation occurs through the receptor-mediated recruitment of coregulator proteins that impact *e*.*g*., chromatin structure *via* promotion of nucleosome remodeling, and TF activity and chromatin accessibility through the covalent modification of proteins (3, 4). Besides the recruitment of coregulators, steroid receptors can influence the chromatin binding and activity of each other as well as other TFs through a variety of crosstalk mechanisms (5, 6). These mechanisms can be direct where the TFs act at the same site, or they can influence each other indirectly through secondary proteins and pathways. Especially in cancers, where TF activity is commonly dysregulated, crosstalk mechanisms play a prominent part in the development, progression, and maintenance of the disease (5, 7–9).

Since the aberrant and persistent AR signaling drives the prostate cancer (PCa) development and progression, the receptor is the main precision target of cancer therapy (5, 10). Besides androgen deprivation therapy (ADT) to limit the availability of activating androgens, new generation of AR antagonizing antiandrogens have been developed to target the sustained AR activation. One such antiandrogen is enzalutamide (ENZ) that is used for the treatment of castration resistant prostate cancer (CRPC) (10). Unfortunately, resistance to ENZ and other antiandrogens frequently prevails within months, leaving CRPC currently incurable. These resistance mechanisms include AR mutations (11) and splice variants (12), as well as gene (13) and enhancer amplification (14), all of which ultimately lead to progressed disease despite the low levels of circulating androgens.

Resistance to antiandrogens can also be attained through steroid receptor crosstalk. In PCa, the AR represses the gene expression of *NR3C1* (gene encoding GR) in treatment-naïve cells (9, 15). In response to the inhibition of AR signaling, *e*.*g*., with ENZ, the expression of *NR3C1* is de-repressed, leading to the elevated levels of GR protein. Subsequently, the upregulated GR replaces the functions of AR and the regulation of its target genes leading to antiandrogen resistance (9, 16). The close similarity between the receptors’ domain structure, consensus binding sequence, and interacting proteins (3, 5) expectedly enable the GR-mediated replacement of AR. The replacement by GR is not complete but is suggested to cover half of AR binding sites (9). In addition to the elevated levels of GR, steroid hormone metabolism also influences the antiandrogen resistance and GR activity through increased cortisol levels upon ENZ treatment (17, 18) as well as glucocorticoid-mediated suppression of adrenal androgen production (19). Moreover, synthetic glucocorticoids such as dexamethasone (Dex), are commonly used as an adjunct therapy to relief side effects of the cancer treatments, likely further stimulating the GR in PCa.

At the clinical level, PCa patients with high GR-positivity showed a poor response to ENZ therapy (9), and increased GR levels were correlated with persistent tumors after androgen synthesis blocker abiraterone treatment (20). Moreover, glucocorticoid usage in combination with ENZ was associated with inferior overall and progression-free survival compared to usage with only ENZ (21). Due to these effects, GR antagonism could seem like an attractive approach to counter this type of antiandrogen resistance. However, GR is ubiquitously expressed and an essential protein for life, which hampers the receptor’s full antagonization. Moreover, CRPC patients treated with ENZ combined with GR antagonist mifepristone did not receive survival benefit over patients treated only with ENZ (22). Since the direct inhibition of GR has not provided the desired outcome, it becomes crucial to investigate the mechanisms of GR signaling in antiandrogen resistance, with the focus on restricting GR signaling in the cancer while retaining the beneficial effects of glucocorticoids at a system-wide level (5).

Most of the current genome-wide knowledge of GR’s transcriptional activity is limited to LREX cells originating from engineered AR overexpressing LNCaP cells that were shaped ENZ-resistant through mouse xenograft (9). In the current study, we explored the genome-wide effects of GR signaling in the same PCa cell background grown with or without ENZ. We show that upregulated GR preferably binds to pre-accessible chromatin sites in ENZ-treated PCa cells, which elucidates the logic behind AR’s partial replacement. Furthermore, we also observe that the action of GR can be restricted using small-molecule inhibitors against coregulators such as p300, advocating their utilization to combat antiandrogen resistance. Finally, we indicate an unexpected role of pioneer factor FOXA1 in the regulation of GR activity.

## MATERIALS AND METHODS

### Cell culture and treatments

VCaP (ATCC, #CRL-2876) cells were maintained in DMEM (Gibco, #41965-039) supplemented with 10% FBS (Cytiva, #11531831) and 1 U/μl penicillin and 1 μg/ml streptomycin (Gibco, #15140122). 22Rv1 (ATCC, #CRL-2505) cells were maintained in RPMI-1640 medium (Gibco, #1187509) supplemented with 10% FBS (Gibco, #11573397), 1 U/μl penicillin, 1 μg/ml streptomycin and 2 mM L-glutamine (Gibco, #A2916801). Enzalutamide-treated cells were cultured with 10 μM enzalutamide (ENZ) (Orion, #ORM-0016678) for at least 21 days before the start of the experiments, passaging or changing the media every 3-4 days. In general, the ENZ treatment time ranged between 21 and 24 days, and the range has been indicated as 3 weeks (3w) in the text and figures. For experiments, the cells were cultured in experiment medium containing 5% charcoal stripped FBS (Gibco, #10270106) for at least two days before the start of hormone treatments. For hormone treatments, 100 nM dexamethasone (Dex) (Sigma-Aldrich, #D4902) and 1 nM or 100 nM 5α-dihydrotestosterone (DHT) (Steraloids, #A2570-000) was used. For experiments with the coregulator inhibitors, the cells were treated with 0.1 μM or 1 μM A-485 p300/CBP HAT (Tocris, #6387), CCS1477 p300/CBP bromodomain (MedChemExpress, #HY-111784), I-BET762 BET bromodomain (Sigma, #SML1272), or BRM014 BRM/BRG1 ATP (MedChemExpress, #HY-119374) inhibitors.

### Antibodies

Primary antibodies: anti-GR (Cell Signaling Technology, #12041), anti-AR (23), anti-FOXA1 (Abcam, #ab23738), anti-TLE3 (Abcam, #ab94972), anti-H3K27ac (Active Motif, #39133), anti-α tubulin (TUB) (Santa Cruz Biotechnology, #sc-5286). Secondary antibodies: goat anti-rabbit (Invitrogen, #G-21234), goat anti-mouse (Zymed Laboratories, #81-6520), donkey anti-rabbit Alexa Fluor 647 (Thermo Fisher, #A-31573).

### Immunoblotting

Proteins were isolated and separated as before (4). The following antibodies were utilized: anti-GR (1:1000), anti-AR (1:10 000), anti-FOXA1 (1:5000), anti-TLE3 (1:1000). An anti-TUB antibody (1:3000) was used as a control for sample loading. Goat anti-rabbit (1:10 000) was used as a secondary antibody for GR, AR, FOXA1 and TLE3, and goat anti-mouse (1:10 000) for tubulin. Protein bands were detected using Pierce ECL Western Blotting Substrate kit (Thermo Fisher, #32106) and Bio-Rad ChemiDoc Imager. Protein level quantifications were calculated using ImageJ software (National Institutes of Health). The samples were first normalized to their TUB loading control, and then to the control ENZ 0 treatment. For ChIP Western blot, samples were first processed as ChIP-seq samples (see below) using 10 μl anti-GR antibody per IP. Subsequently, instead of reverse crosslinking and elution, input and IP samples were incubated at 95 °C in SDS buffer. Next, the samples were immunoblotted, detected, and quantified as indicated above.

### Immunostaining of GR, confocal imaging, and image analysis

The ±ENZ-treated cells (VCaP: 100 000 per well; 22Rv1: 40 000 per well) were cultured in charcoal stripped medium on 8-well chamber slides (Ibidi GmbH, #80826). Before staining, the cells were treated with EtOH or 100 nM Dex for 1 h at 37 °C. For fluorescent staining, the cells were fixed with 4% paraformaldehyde in 0.1 M sodium phosphate buffer, pH 7.4 (PB) for 20 min, and washed with PB. The fixed cells were permeabilized for 15 minutes with 0.1% Triton X-100 and 1% BSA, blocked with 1% BSA for 20 minutes, and incubated o/n at 4 °C with anti-GR antibody (1:500). After washing with PB, the cells were incubated for two hours with Alexa Fluor® 647 labeled secondary antibody (1:500), and nuclei were labeled with 4’,6-diamidino-2-phenylindole (DAPI, 1 μg/mL) (Sigma-Aldrich, #D8417). The fluorescent images were obtained with a Zeiss Axio Observer inverted microscope (40 × NA 1.3 oil objective) equipped with a Zeiss LSM 800 confocal module (Carl Zeiss Microimaging GmbH). Image processing was performed using ZEN 2.5 (blue) software (Carl Zeiss Microimaging GmbH), and the images were quantified with ImageJ software (National Institutes of Health).

### RNA extraction and RT-qPCR

For RNA isolation, the cells were seeded onto 6-well plates, 1 million ±ENZ VCaP cells and 300 000 ±ENZ 22Rv1 cells per well, respectively. For RNA interference, the cells were reverse transfected with siRNA using Opti-MEM (Thermo Fisher, #31985070) and Lipofectamine RNAiMAX Transfection Reagent (Invitrogen, #13778150) according to manufacturer’s instructions. The following siRNA SMARTpools (Horizon/Dharmacon) at 60-80 nM concentration were used: ON-TARGETplus Non-targeting Control Pool (siNON, #D-001810-10), ON-TARGETplus Human NR3C1 siRNA (siNR3C1, L-003424-00-0020), ON-TARGETplus Human FOXA1 siRNA (siFOXA1, #L-010319-00-0050) or ON-TARGETplus Human TLE3 siRNA (siTLE3, #L-019929-00-0020). Medium was changed into charcoal stripped medium (±ENZ) three days before the isolation. The effectiveness of each siRNA was validated with immunoblotting as described above. For experiments with coregulator inhibitors, the cells were treated with an inhibitor or DMSO for 24 h before extraction of RNA. The cells were treated with vehicle or steroid hormone 18 h prior to the RNA extraction. Total RNA was extracted using TriPure Isolation Reagent (Roche, #1166715700). Quality and quantity of the isolated RNA were determined with NanoDrop One (Thermo Scientific). 1 μg of total RNA from each sample was converted into cDNA with First Strand cDNA Synthesis Kit (Roche, #04897030001). Primers used in the RT-qPCR are listed in Supplementary Table S1. The qPCR was performed using LightCycler 480 Instrument II (Roche) and the relative gene expression was calculated using the 2^-(ΔΔCt)^ method, where ΔΔC_t_ is ΔC_t(treatment)_-ΔC_t(vehicle)_ and ΔC_t_ is C_t(target gene)_-C_t(control gene)_. *RPL13A* was used as a control gene. P-values were calculated using One-way ANOVA with Bonferroni *post hoc* test.

### RNA-seq and data analysis

Samples were treated as indicated above and RNA was extracted with RNeasy Plus Mini kit (Qiagen, #74134) after which the RNA integrity number (RIN) was measured with Agilent 2100 Bioanalyzer using RNA 6000 Nano kit (Agilent, #5067-1511). All samples had RIN >9, indicating high integrity of RNA. RNA-sequencing (RNA-seq) libraries were generated using NEBNext Poly(A) mRNA Magnetic Isolation Module (New England BioLabs, #E7490) and NEBNext Ultra II Directional RNA Library Prep Kit (New England BioLabs, #E7765) according to manufacturer’s protocol. Library quality was assessed with Agilent 2100 Bioanalyzer using DNA 1000 Analysis kit (Agilent, #5067-1504). Two to three biological replicate samples were sequenced with Illumina NextSeq 500 (75SE). RNA-seq data was aligned to hg38 genome using STAR2.7 (24) with default settings and max 10 mismatches and max 10 multi-mapped reads. Differentially expressed genes were then analyzed with DESeq2 (25) through HOMER (26) for all comparisons. Total count per gene was calculated using transcripts per million (TPM) normalization and other than protein coding genes were filtered out as outliers. Genes with TPM >0.5 at least in one sample in any treatment were considered as expressed. Differentially expressed genes were then defined as FDR <0.05 and log2 fold change ±0.5 between differently treated samples. All RNA-seq data values are presented in Supplementary Table S2. Heatmap displaying Z-scores for each replicate in each condition were drawn using heatmap.2 in gplots R package (R project). Differentially expressed gene clusters were used in gene set enrichment analysis (GSEA) (27) or Metascape (28) pathway analysis with default settings. For GSEA, hallmark and biological process ontology gene sets were utilized. Prediction of transcriptional regulators from differentially expressed genes, epigenetic landscape *in silico* deletion analysis (LISA) (29) was used with default settings. For coregulator enrichment analysis, TcoFBase (30) TcoF gene set enrichment was used with Dex-regulated genes using FDR adjustment with default setting. The Cancer Genome Atlas (TCGA) data was obtained from cBioPortal (31) and Xena browser (32). The Cancer Cell Line Encyclopedia (CCLE) data was obtained from Expression Atlas (33) and supplemental material (34). GSEA, Metascape, LISA and TcoFBase pathway enrichment data is presented in Supplementary Table S3. Publicly used RNA-seq and proteomics data is shown in Supplementary Table S4.

### ChIP-seq

For chromatin immunoprecipitation (ChIP), VCaP cells were seeded onto 10 cm plates, 10 million untreated and 11 million ENZ-treated cells per plate. 22Rv1 cells were seeded 3 million untreated and 4 million ENZ-treated cells per plate. For RNA interference, VCaP cells were treated with 80 nM of siNON or siFOXA1 as described above. The effectiveness of FOXA1 depletion was validated with immunoblotting as described above. ChIP-seq experiments were done as previously described (35). Briefly, the cells were treated with vehicle, Dex (100 nM) or DHT (1 nM or 100 nM) for 1 h prior to the start of ChIP. Chromatin was fragmented to an average size of 150–300 bp by sonication (Diagenode, #UCD-300). Antibodies were coupled to magnetic protein G Dynabeads (Invitrogen, 10004D) for at least 16 h, and sonicated lysates were incubated with antibody-coupled beads for o/n at 4 °C. Antibodies used per IP: GR, 12.5 μl; AR, 3 μl; FOXA1, 2 μg; TLE3, 2 μg; H3K27ac, 2 μg. Two to four IP samples were pooled for one ChIP-seq sample. Sequencing libraries were generated using NEBNext Ultra II DNA Library Prep Kit (New England BioLabs, #E7645L) according to manufacturer’s protocol. Analysis of library quality was done with Agilent 2100 Bioanalyzer using DNA 1000 Analysis kit (Agilent, #5067-1504). Two biological replicate samples were sequenced with Illumina NextSeq 500 (75SE).

### ATAC-seq

For assay for transposase-accessible chromatin with sequencing (ATAC-seq), VCaP cells were seeded onto 10 cm plates, 10 million untreated and 11 million ENZ-treated cells per plate, and 22Rv1 cells were seeded 3 million untreated and 4 million ENZ-treated cells per plate. ATAC-seq experiments were done as previously described (36, 37). Briefly, the cells were treated with vehicle, 100 nM Dex or 1 nM DHT for 1 h prior to the start of nuclei isolation. For nuclei isolation, the cell pellets were resuspended in a concentration of 5 million cells per ml in Buffer A [15 mM Tris–HCl (pH 8), 15 mM NaCl, 60 mM KCl, 1 mM EDTA, 0.5 mM EGTA, 0.5 mM spermidine (Sigma-Aldrich, S2626), 1X protease inhibitor cocktail]. Subsequently, equal volume of Buffer A with 0.04% (w/v) IGEPAL (Sigma, #I8896) was added, to obtain a concentration of 2.5 million cells per ml with 0.02% (w/v) IGEPAL. Samples were incubated on ice for 10 min, washed once with Buffer A without IGEPAL, and two times with ATAC resuspension buffer (10 mM NaCl, 10 mM Tris-HCl, 3 mM MgCl_2_). Isolation of nuclei was verified by Trypan Blue counting. Subsequently, 100 000 nuclei were subjected to Tn5 transposition reaction using 2.5 μl TDE1 from Nextera DNA Library Prep Kit (Illumina, #FC-121-1030). After adding the transposition reaction mix, the samples were incubated 45 min at 37 °C with 800 rpm shaking, and subsequently DNA was purified using Monarch PCR & DNA Cleanup Kit (New England BioLabs, #T1030). DNA fragments were amplified with PCR and samples were barcoded using published primers (36). Amplified fragments were size selected (150 bp-800 bp) using SPRIselect beads (Beckman Coulter, #B23318). Analysis of library quality was done with Agilent 2100 Bioanalyzer using High Sensitivity DNA Analysis kit (Agilent, #5067-4626). Two biological replicate samples were sequenced with Illumina NextSeq 500 (40PE).

### ChIP-seq and ATAC-seq data analysis

For ChIP-seq, read quality filtering and alignment to hg38 genome using Bowtie (38) was performed as previously described (35). For ATAC-seq, read quality filtering and alignment to hg38 genome using Bowtie2 (39) was performed as previously described (37). Downstream data analysis was performed using HOMER (26). Peaks were called using default parameters on findPeaks with style factor, FDR <0.01, >25 tags, >4-fold over control sample and local background. ChIP input sample was used as a control. DESeq2 (25) through getDiffrentialPeaksReplicates.pl was used to isolate differential binding peaks (FDR <0.1, fold change >2) between specified treatments (unchanged, UN; increased, UP; decrease, DN). ChIP peaks are presented in Supplementary Table S5. Aggregate plots and heatmaps were generated with 10 bp or 20 bp bins surrounding ±1 kb area around the center of the peak. All plots were normalized to 10 million mapped reads and further to local tag density, tags per bp per site, whereas box plots and scatter plots represented log2 tag counts. Statistical significance in the box plots was determined using One-way ANOVA with Bonferroni *post hoc* test. Box plots were generated using Tukey method with interquartile range (IQR) depicting the 25th, 50th and 75th percentile as box with the median as black bar. The whiskers extend 1.5xIQR beyond the box, and outliers are not depicted but are included in the statistical analyses. *De novo* motif searches were performed using findMotifsGenome.pl with the following parameters: 200 bp peak size window, strings with 2 mismatches, binomial distribution to score motif p-values, and 50 000 background regions. Enriched motifs are presented in Supplementary Table S6. Gene-to-peak association was performed with annotatePeaks.pl measuring linear distance from target gene TSS to center of the peak. Distances were shown as cumulative distribution function from 0 to 100 kb with 10 kb intervals. Statistical significance was calculated with Kolmogorov–Smirnov test comparing differentially expressed Dex-regulated genes to expressed genes. Replacement of AR by GR was measured by direct overlap of the receptor binding sites and presented as Venn diagrams. Pre-accessible and *de novo* GR binding sites were defined based on ATAC-seq or DNase-seq data. *De novo* sites had less than 1 log2 tags and pre-accessible sites had more than 1 log2 tags in untreated condition. PCa patient ATAC-seq and FOXA1 ChIP-seq data tag counts at GR binding sites were calculated with annotatePeaks.pl and heatmaps were generated using hierarchical clustering with Euclidean distance. Enrichment of binding sites to GO biological processes, GREAT (40) was used with basal plus extension association. GREAT pathway enrichment data is presented in Supplementary Table S3. Publicly used ChIP-seq, ATAC-seq and DNase-seq data is shown in Supplementary Table S4.

### Cell proliferation (MTS) assay

For MTS assay, the cells were seeded onto 96-well plates: 30 000 VCaP cells and 10 000 22Rv1 cells per well in charcoal-stripped medium (±ENZ). For RNA interference, the cells were transfected with siNON or siFOXA1 as described earlier. Wells containing only medium were used to subtract background. The start of the treatments was designated as day 0 (d0). The cell proliferation was assessed at d0, d1, d2, d3 and d4 using colorimetric CellTiter 96 Aqueous One Solution Cell Proliferation Assay (MTS) kit (Promega, #G3580) according to manufacturer’s protocol. The absorbance was measured at 492 nm wavelength using Multiskan EX plate reader (Thermo Scientific). Results were normalized to d0 and shown as fold change. P-values were calculated using Two-way ANOVA with Bonferroni *post hoc* test.

## RESULTS

### GR regulates AR associated pathways in antiandrogen-treated prostate cancer cells

To investigate the early impact of GR-mediated antiandrogen resistance through the direct replacement of androgen signaling (9), we utilized two PCa cell lines, VCaP and 22Rv1, with the former representing androgen-dependent and the latter androgen-independent cells (16, 41). Basally, AR displays higher protein levels in VCaP compared to 22Rv1 cells, though more putative AR splice variants are expressed in the latter cells (Supplementary Figure S1A). In comparison, the basal protein levels of GR are substantially higher in 22Rv1 compared to VCaP cells. To de-repress the AR-mediated inhibition of *NR3C1* (gene encoding GR) expression (5, 15), the VCaP and 22Rv1 cells were grown in the presence of 10 μM enzalutamide (ENZ) for at least three weeks (ENZ 3w) (see Materials and Methods for details). Immunofluorescence analysis indicated that the mean nuclear intensity of Dex-activated GR significantly increased after ENZ 3w in both VCaP and 22Rv1 cells (Figure 1A-B, Supplementary Figure S1B-C). In addition, GR protein levels were increased (Supplementary Figure S1A) to a similar degree as has been shown by others (17, 18, 42). Finally, RNA-seq analyses of untreated (ENZ 0) and ENZ 3w treated VCaP and 22Rv1 cells verified the significant increase in *NR3C1* transcript levels upon ENZ 3w (Figure 1C). This is complementary to the RNA-seq analysis of *NR3C1* transcript levels before and after androgen deprivation and/or ENZ treatment from PCa patients and mouse xenografts (Supplementary Figure S1D-H). To verify that AR signaling is inhibited, we performed RNA-seq, AR ChIP-seq and ATAC-seq in ENZ 0 and ENZ 3w treated VCaP and 22Rv1 cells. In both cell lines, the DHT-regulated transcriptome, AR chromatin binding and AR-induced chromatin accessibility was drastically inhibited by ENZ 3w (Supplementary Figure S1I-L). Next, we focused on the GR-regulated transcriptome in ENZ-treated cells.

**Figure 1.**
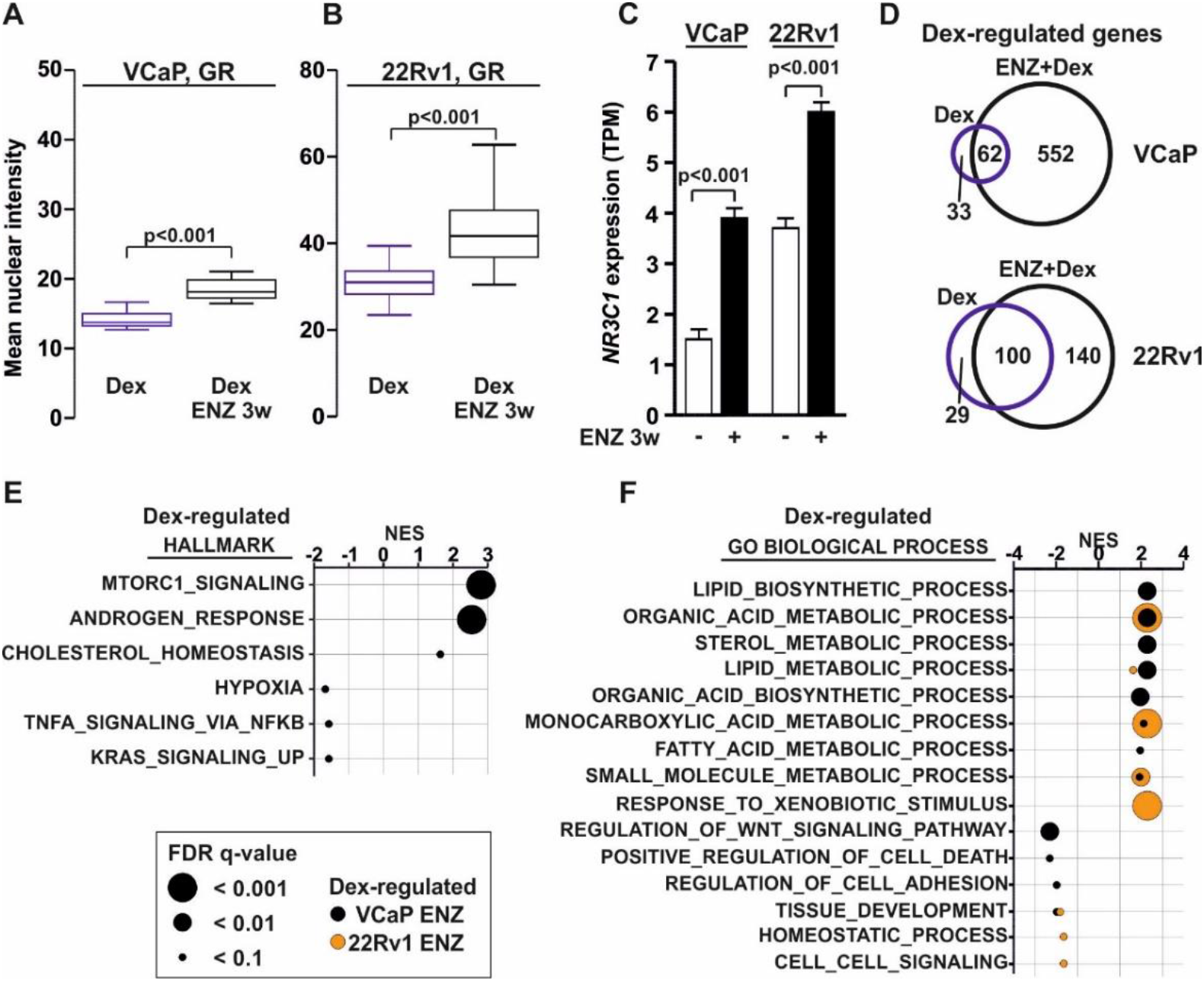
GR-regulated transcriptome is associated with AR pathways in antiandrogen-treated prostate cancer cells. (A-B) Box plots represent quantification of GR immunofluorescence (IF) in Dex-treated VCaP (A) or 22Rv1 (B) ENZ 0 or ENZ 3w cells. Data shown as mean nuclear intensity, VCaP: n=15 images; 22Rv1: n=30 images. Statistical significance calculated with unpaired t-test. (C) Bar graphs depict *NR3C1* expression levels in VCaP and 22Rv1 ENZ 0 or ENZ 3w cells from RNA-seq data. Data shown as transcripts per million (TPM). Statistical significance calculated with unpaired t-test. (D) Venn diagrams of Dex-regulated genes in VCaP and 22Rv1 cells from RNA-seq data after Dex (blue circle) or ENZ+Dex (black circle) treatment. (E-F) Gene set enrichment analysis (GSEA) of Dex-regulated genes from VCaP (black) or 22Rv1 (orange) ENZ 3w cells at hallmark (E) or GO biological process (F) pathways. Data shown as normalized enrichment score (NES) with circle size depicting FDR q-value.

RNA-seq analyses indicated that Dex-regulated transcriptome is vastly expanded in ENZ 3w cells (Figure 1D, Supplementary S2A-B, Supplementary Table S2). Complementary to the basal GR protein levels, without ENZ treatment 22Rv1 cells harbored more Dex-regulated genes than VCaP cells. However, VCaP cells displayed a more extensive increase in the number of Dex-regulated genes upon ENZ 3w. The majority of the Dex-regulated genes after ENZ 3w were cell line-selective (Supplementary Figure S2C). As an example, *FKBP5* and *KLK3* (gene encoding PSA) were similarly Dex-regulated regardless of ENZ treatment whereas *SGK1* and *KLF9* were more efficiently Dex-regulated in ENZ 3w cells (Supplementary Figure S3A-B). Using control siRNA (siNON) and siRNA against *NR3C1* (siNR3C1), we verified that the Dex-regulation of *SGK1* and *KLF9* was dependent on GR (Supplementary Figure S3C-E). In addition, one-day ENZ withdrawal after ENZ 3w reduced *NR3C1* expression and restored DHT-regulation of *FKBP5* and *KLK3* (Supplementary Figure S3F). To investigate if the Dex-regulated genes in ENZ-treated cells were associated with similar pathways as DHT-regulated genes in ENZ 0 cells, we performed gene set enrichment analyses (GSEA). As expected, the DHT-regulated genes in ENZ 0 cells were positively associated with androgen response as well as several lipid and steroid metabolic pathways, and upon ENZ 3w these pathways presented negative association (Supplementary Figure S2D-E, Supplementary Table S3). Interestingly, like DHT-regulated genes in ENZ 0 cells, the Dex-regulated genes were positively associated with the androgen response in ENZ 3w VCaP cells (Figure 1E). In addition, the Dex-regulated genes in ENZ 3w VCaP and 22Rv1 cells were positively associated with multiple lipid and steroid metabolic pathways (Figure 1F). Dex-regulated genes exhibited similar pathway enrichment with Metascape analyses (Supplementary Figure S2F, Supplementary Table S3). Although similar effects have been shown with a restricted number of target genes (9), our results indicated that on a genome-wide transcriptome level and in the same cellular background, GR-regulated genes in ENZ-treated PCa cells are enriched at biological pathways normally regulated by AR.

### Chromatin binding of GR occurs at pre-accessible sites

To complement the RNA-seq analyses, we performed GR ChIP-seq from VCaP and 22Rv1 cells using the same ENZ treatment conditions. We defined the GR binding sites (GRBs) as unchanged (ENZ-UN) and increased (ENZ-UP) depending on their response to ENZ 3w treatment (see Materials and Methods for details). Significantly decreased GRBs upon ENZ 3w were not found in the cells. In agreement with transcriptome analyses, ENZ 3w expanded GRBs in both cell lines (Figure 2A, Supplementary Figure S4A). 22Rv1 cells displayed more GR chromatin binding without ENZ treatment, whereas VCaP cells exhibited a more substantial increase in the number of GRBs after ENZ 3w (Figure 2C-D, Supplementary Figure S4B-C, Supplementary Table S5). The latter was mirrored by the increase in GR protein levels on chromatin in VCaP cells (Figure 2B). Like ENZ-UP, also the ENZ-UN sites showed increased GR chromatin binding when tag density was quantified (Figure 2D, Supplementary Figure S4C). Association of Dex-regulated genes to GRBs indicated that in VCaP cells, only ENZ-UP sites were significantly linked with Dex-regulated genes observed in the ENZ 3w cells (shared or ENZ gene clusters) (Figure 2E). In direct comparison, ENZ-UN sites in 22Rv1 cells were linked with Dex-regulated genes observed in ENZ 3w cells (shared or ENZ gene clusters) (Supplementary Figure S4D). Thus, the transcriptome and GR cistrome analyses suggested that in ENZ 3w cells, GR operates through new binding sites in androgen-dependent VCaP cells while in androgen-independent 22Rv1 cells GR utilizes existing binding sites.

**Figure 2.**
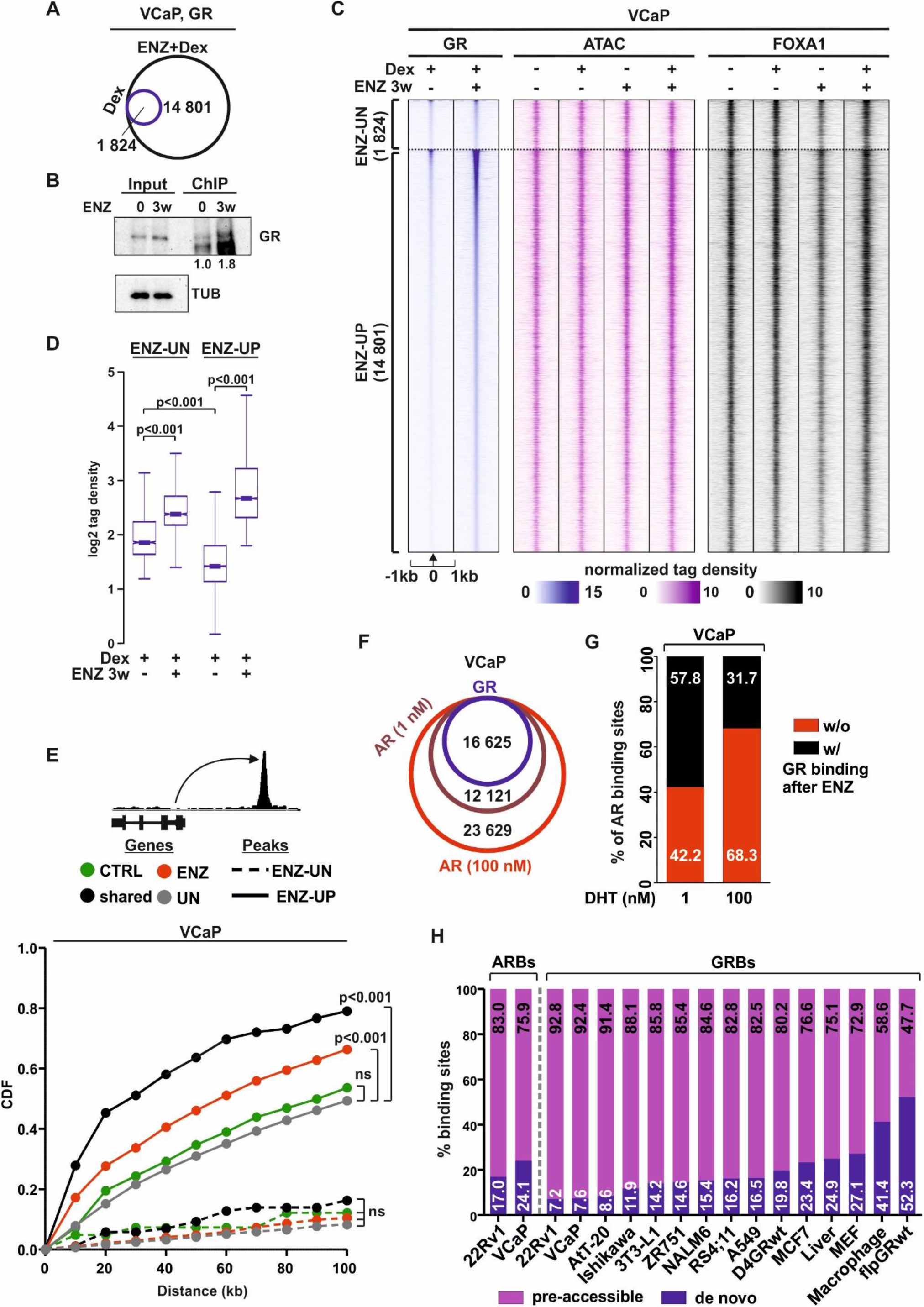
GR preferably binds to pre-accessible chromatin sites in prostate cancer cells. (A) Venn diagrams of GR chromatin binding in VCaP cells from GR ChIP-seq after Dex (blue circle) or ENZ+Dex (black circle) treatment. (B) Immunoblotting of GR from GR ChIP samples VCaP ENZ 0 and ENZ 3w cells. Immunoblotting of GR and TUB from ChIP input used as control. Quantification of GR chromatin levels to input is shown below the immunoblot. (C) GR ChIP-seq, ATAC-seq, and FOXA1 ChIP-seq profiles at ENZ-UN and ENZ-UP sites in VCaP ENZ 0 and ENZ 3w cells. ENZ-UN represents unchanged and ENZ-UP increased GR binding sites. Each heatmap represents ±1 kb around the center of the GR peak. Binding intensity (tags per bp per site) scale is noted below on a linear scale. (D) Box plots represent the normalized log2 tag density of GR ChIP-seq at ENZ-UN and ENZ-UP sites. Statistical significance calculated using One-way ANOVA with Bonferroni *post hoc* test. (E) Association of Dex-regulated genes in VCaP ENZ 0 and ENZ 3w cells to ENZ-UN and ENZ-UP GR binding sites. Different subsets of Dex-regulated genes are color coded, and ENZ-UN depicted as dotted line and ENZ-UP as solid line. Statistical significance calculated with Kolmogorov–Smirnov test. (F) Venn diagrams of GR and AR chromatin binding in VCaP cells from GR ChIP-seq (blue circle) and AR ChIP-seq treated with 1 nM (magenta circle) or 100 nM (red circle) of DHT. (G) Bar graphs depict percentage overlap of AR binding sites with (black) or without (red) GR binding. (H) Bar graphs depict percentage overlap of AR binding sites (ARBs) or GR binding sites (GRBs) with pre-accessible (purple) or *de novo* (blue) chromatin sites. Cell lines or tissues analyzed are indicated below the bar graphs. All heatmaps and box plots are normalized to a total of 10 million reads.

To analyze the replacement of AR binding sites (ARBs) by GR in ENZ-treated cells, we measured the direct overlap of ARBs in ENZ 0 cells versus GRBs in ENZ 3w cells. The degree of ARB replacement by GR binding was around 58% and 76% in VCaP and 22Rv1 cells, respectively, when AR was activated with 1 nM of DHT (Figure 2F-G, Supplementary Figure S4E-G). This concentration of DHT was also used in LREX cells displaying 46% overlap between ARBs and GRBs (Supplementary Figure S4F-G) (9). However, when the VCaP and 22Rv1 cells were treated with hundred-fold more (100 nM) DHT, the replacement of ARBs by GR decreased to around 32% and 20%, respectively. This reflects the increased number of ARBs gained from the higher DHT concentration. Thus, the data suggests that the partial replacement of AR by GR occurs primarily at ARBs which are sensitive to low levels of androgens.

Intriguingly, it has been shown that GRBs sensitive to Dex concentration occur at pre-accessible chromatin sites. These sites are commonly characterized by the presence of active histone marks and chromatin being open prior to the hormone stimulus (43). In comparison, these events are not present prior to GR binding at closed chromatin (*de novo*) sites. Due to this, we performed ATAC-seq from VCaP and 22Rv1 cells to analyze the chromatin accessibility at GRBs. While chromatin accessibility was significantly increased after Dex treatment in ENZ 3w VCaP cells and in 22Rv1 cells regardless of ENZ treatment (Supplementary Figure S5A-B), GRBs occurred at pre-accessible chromatin sites (Figure 2C, Supplementary Figure S4B). Furthermore, sorting of VCaP ARBs based on chromatin pre-accessibility, illustrated that while AR can bind to *de novo* sites, GRBs overlap only with the most pre-accessible ARBs (Supplementary Figure S5C). In support of binding to open chromatin sites, GRBs displayed the basal enrichment of active histone marks, H3K4me1 and H3K27ac, in both VCaP and 22Rv1 cells (Supplementary Figure S5D). Moreover, H3K27ac ChIP-seq from 22Rv1 cells indicated that ENZ 3w cells retained the enrichment of H3K27ac and Dex treatment increased the enrichment more clearly in ENZ 3w compared to ENZ 0 cells (Supplementary Figure S5E). Finally, GRBs in VCaP and 22Rv1 cells presented mostly open chromatin configuration in the PCa patients (Supplementary Figure 5F-G). The example loci of GR ChIP-seq and ATAC-seq from VCaP and 22Rv1 cells are shown in Supplementary Figure S6A-D. To analyze if GRBs’ occurrence at pre-accessible sites is specific for PCa cells, we obtained GR ChIP-seq and ATAC-seq or DNase-seq datasets from other cell lines and tissues (Supplementary Table S4). Subsequently, we divided GRBs based on chromatin accessibility to pre-accessible or *de novo* sites (see Materials and Methods for details). Interestingly, over 90% of GRBs in VCaP and 22Rv1 cells occurred at pre-accessible sites, the highest percentage of all analyzed cells (Figure 2H). Only mouse pituitary tumor AtT-20 cells showed over 90% pre-accessibility at GRBs. Contrastingly, in A549 lung cancer cell line and mouse liver tissue, which represent a more common cellular background to GR, GRBs displayed similar degrees of pre-accessible enrichment as ARBs in 22Rv1 and VCaP cells. Thus, the binding of GR in PCa cells is specifically restricted to chromatin sites that are already open prior to hormone and ENZ treatments. This suggests that correspondingly the replacement of ARBs by GR would be limited by the chromatin accessibility.

### Inhibition of p300 enzymatic activity restricts cell proliferation and GR’s transcriptional activity

Since the regulation of chromatin accessibility and enhancer activation is tightly dependent on coregulators recruited to chromatin by TFs (44, 45), we hypothesized that through the inhibition of coregulatory activity, GR’s transcriptional regulation would be inhibited. To narrow down the potential coregulators that influence the action of GR, we utilized TcoF gene set enrichment (30). The analyses indicated that BET family members, EP300/CREBBP (p300/CBP), SWI/SNF complex members, and MED1 were enriched at GR-regulated target genes in VCaP and 22Rv1 cells (Supplementary Figure S7A, Supplementary Table S3). Since chromatin accessibility was significantly higher at ARBs shared with GR compared to ARBs without GR in VCaP and 22Rv1 cells (Supplementary Figure S7B-C), we expected that coregulators prominently involved at GR-regulated enhancers should display similar difference. We utilized H3K27ac, p300/CBP-mediated histone modification, as proxy for p300 occurrence. Interestingly, ChIP-seq analyses from VCaP cells indicated that H3K27ac and SWI/SNF ATPase SMARCA4 (BRG1) displayed significantly more enrichment at GR-bound ARBs than at ARBs without GR occupancy (Supplementary Figure S7B). No difference was observed with BRD4 and MED1 binding to these sites. In 22Rv1 cells, H3K27ac as well as BRD4 displayed significantly more enrichment at ARBs with GR than ARBs without GR (Supplementary Figure S7C). These results suggest that BRD4, BRG1, and especially p300/CBP could influence GR’s transcriptional action in PCa cells. To test this, we utilized several small chemical inhibitors targeting p300/CBP HAT (A-485) (46), p300/CBP bromo (CCS1477) (47), BET bromo (I-BET762) (48), and BRM/BRG1 ATP domain (BRM014) (49) (Figure 3A). Gene expression analysis of several model target genes of GR indicated that overall, p300/CBP inhibition with A-485 or CCS1477 had the most prominent inhibitory effect in both VCaP and 22Rv1 cells treated with or without ENZ (Figure 3B-E, Supplementary Figure S8A-D). Furthermore, p300/CBP HAT domain inhibitor A-485 was the most effective compound in restricting VCaP and 22Rv1 cell proliferation (Figure 3F-G). This was especially prominent in ENZ 3w cells. Taken together, the transcriptional activity of GR in antiandrogen-treated PCa cells can be efficiently restricted with the inhibition of p300/CBP activity.

**Figure 3.**
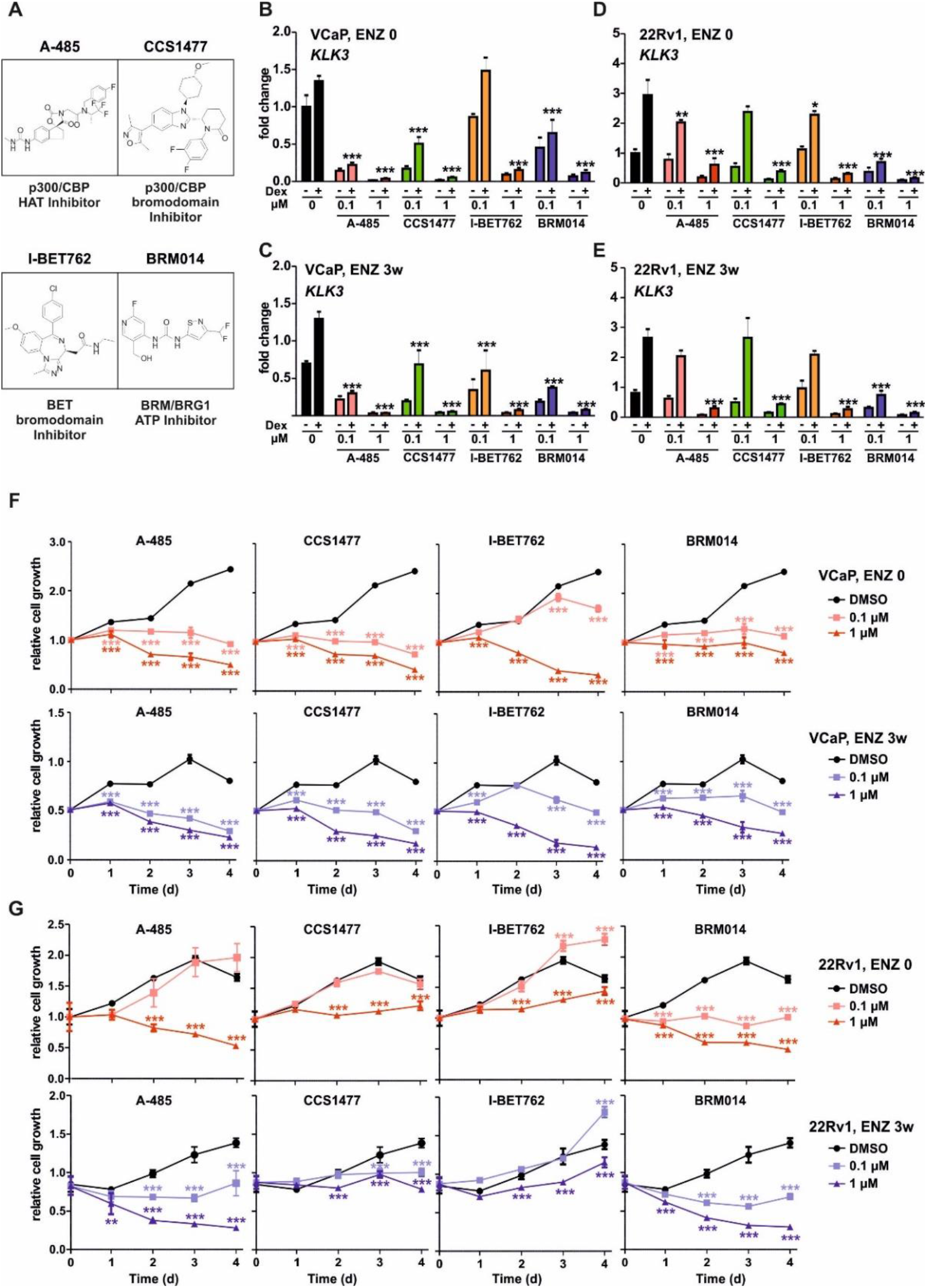
p300 enzymatic activity inhibition hinders cell proliferation and GR transcriptional regulation. (A) Depiction of chemical structures of coregulator inhibitors. A-485, p300/CBP histone acetyl transferase (HAT) domain inhibitor; CCS1477, p300/CBP bromodomain inhibitor; I-BET762, bromodomain and extra terminal domain (BET) bromodomain inhibitor; BRM014, BRM/BRG1 ATP domain inhibitor. (B-E) Bar graphs depict *KLK3* gene expression analysis in VCaP ENZ 0 (B), VCaP ENZ 3w (C), 22Rv1 ENZ 0 (D), and 22Rv1 ENZ 3w (E) cells treated with or without Dex in the presence or absence of 0.1 or 1 μM of indicated inhibitor. Bars represent mean ±SD, n=3. Statistical significance calculated using One-way ANOVA with Bonferroni *post hoc* test. (F-G) Line graphs depict change in cell proliferation as function of time in VCaP (F) and 22Rv1 (G) ENZ 0 and ENZ 3w cells with or without indicated chemical inhibitor treatment. Each data point represents mean ±SD, n=3. Statistical significance calculated using Two-way ANOVA with Bonferroni *post hoc* test. *, p<0.05; **, p<0.01; ***, p<0.001.

### Depletion of FOXA1 potentiates chromatin binding of GR

Besides coregulators, several TFs are known to influence the transcriptional action of GR (2, 5). In support of this, the specific enrichment of GRBs to pre-accessible chromatin suggests that other TFs likely influence the receptor chromatin occupancy. To investigate this, we performed *de novo* motif analyses and discovered that the most prevalent motif at GRBs was the sequence for pioneer factor FOXA1 (Supplementary Figure S9A, Supplementary Table S6). Furthermore, the Dex-regulated genes through LISA (29) predicted FOXA1 to be one of the most crucial factors in the ENZ-treated cells alongside with AR and GR (Supplementary Figure S9B). FOXA1 ChIP-seq from VCaP and 22Rv1 cells indicated that FOXA1 chromatin occupancy is evident at GRBs (Figure 2C, Supplementary Figure S4B, S9C-D). Moreover, GRBs in VCaP and 22Rv1 cells displayed FOXA1 chromatin binding in PCa patients (Supplementary Figure S9E-F). These data suggest FOXA1 could have a prominent role in GR binding in PCa cells.

FOXA1 is known to modulate the chromatin binding of AR (50, 51), as well as GR (51). However, the latter has only been shown in engineered LNCaP cells possessing integrated rat GR. To assess the FOXA1’s prominent role in modulating GR action in native PCa cells, we depleted FOXA1 from VCaP cells using siRNA (siFOXA1) (Supplementary Figure S10A). GR ChIP-seq analyses indicated that the siFOXA1 increased GR chromatin binding at both ENZ-UN and ENZ-UP sites (Figure 4A-B). Intriguingly, cross-comparison of the siFOXA1 versus siNON in ENZ 0 cells and that of the siNON in ENZ 3w versus ENZ 0 cells, showed strong positive correlation in the GR chromatin occupancy (Figure 4C). The correlation was absent if the latter was cross-compared with siFOXA1 versus siNON in ENZ 3w cells (Figure 4D). Thus, the potentiation of GR chromatin binding can be achieved either with ENZ treatment or with the depletion of FOXA1. However, ENZ-UP sites presented significantly increased GR tag density in the ENZ 3w cells after siFOXA1 (Figure 4A). Further analysis of ENZ-UP sites indicated that while the majority of GRBs were unresponsive to siFOXA1 (siFOXA1-UN), some GRBs were lost (siFOXA1-DN), and some gained (siFOXA1-UP) (Supplementary Figure S10B-C). Furthermore, new GRBs were observed after both ENZ 3w and siFOXA1 (*De novo*, siFOXA1-UP). However, ENZ-UP siFOXA1-UP and *de novo* siFOXA1-UP GRBs displayed only modest binding of FOXA1, H3K4me1 and H3K27ac (Supplementary Figure S10C-D), and high enrichment of GRE sequence (Supplementary Figure S10E). This suggests that FOXA1 does not actively inhibit GR binding to FOXA1 binding sites. GREAT analyses (40) indicated that ENZ-UP siFOXA1-UP and *de novo* siFOXA1-UP GRBs were related to the metabolic processes and regulation of cell division pathways (Supplementary Figure S10F). In the case for the latter, the depletion of FOXA1 has been linked to increased PCa cell proliferation (50). In agreement with this, a significant increase in cell proliferation was measured after siFOXA1 in Dex-treated ENZ 0 and ENZ 3w cells (Figure 4E). Thus, FOXA1 seems to indirectly restrict GR action and curb glucocorticoid-induced cell proliferation.

**Figure 4.**
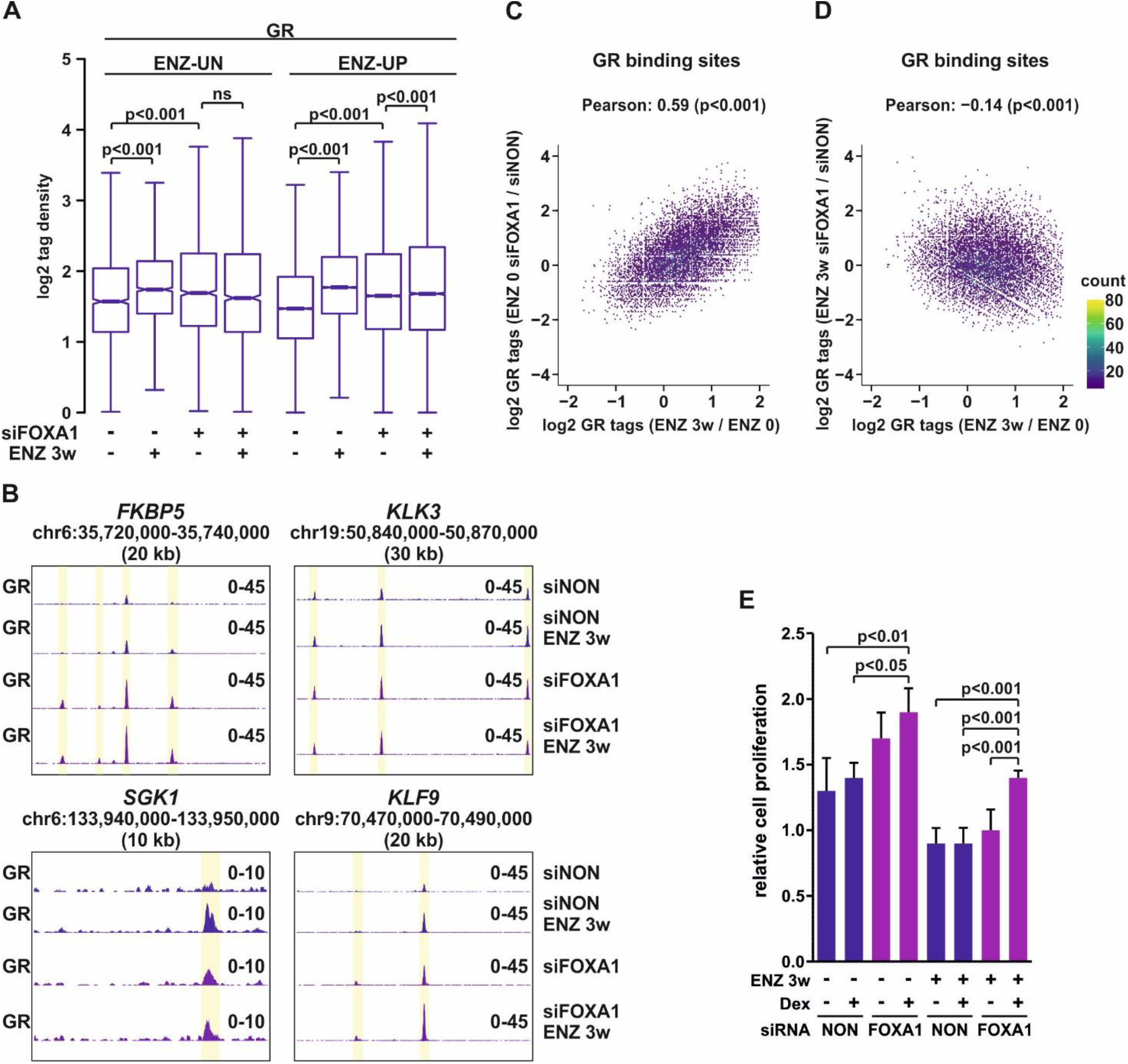
GR chromatin binding is potentiated upon FOXA1 depletion. (A) Box plots represent log2 tag density of GR ChIP-seq at ENZ-UN and ENZ-UP sites in VCaP ENZ 0 and ENZ 3w cells with or without siFOXA1 treatment. Statistical significance calculated using One-way ANOVA with Bonferroni *post hoc* test. (B) Genome browser track examples of GR ChIP-seq at *FKBP5, KLK3, SGK1*, and *KLF9* loci in VCaP ENZ 0 and ENZ 3w cells treated with siNON or siFOXA1. (C-D) Scatter plots of GR ChIP-seq log2 tag density of ENZ 3w/ENZ 0 (x-axis) and ENZ 0 siFOXA1/ENZ 0 siNON (y-axis) (C), and ENZ 3w/ENZ 0 (x-axis) and ENZ 3w siFOXA1/ENZ 3w siNON (y-axis) (D). (E) Bar graphs depict relative cell proliferation in VCaP ENZ 0 and ENZ 3w cells treated with or without Dex in the presence or absence of siFOXA1 for 4 days. Data is normalized to cell proliferation at day 0, and represents mean ±SD, n=3-6. Statistical significance calculated using One-way ANOVA with Bonferroni *post hoc* test. All genome browser tracks, scatter and box plots are normalized to a total of 10 million reads.

### FOXA1 represses *NR3C1* gene expression *via* corepressor TLE3

Since GR chromatin binding manifested a global increase upon FOXA1 depletion, it is plausible that FOXA1 is involved in the repression of *NR3C1* expression. The AR-mediated repression of *NR3C1* has been postulated to occur through EZH2 (15) and the locus contains one prominent ARE-containing ARB displaying the enrichment of EZH2 (Supplementary Figure S11A). In comparison, the two FOXA1 binding sites located at the *NR3C1* promoter show no EZH2 enrichment, suggesting that FOXA1-mediated repression occurs through alternative mechanisms. Considering that FOXA1 is known to recruit the corepressor TLE3 to chromatin (52) and that TLE3 has been implicated in the repression of *NR3C1* expression in PCa cells (53), we hypothesized that FOXA1-mediated repression could operate through TLE3. Interestingly, the FOXA1 binding sites at the *NR3C1* promoter exhibited binding of TLE3 in LNCaP, 22Rv1, and VCaP cells (Figure 5A). Moreover, TLE3 ChIP-seq analyses indicated that depletion of FOXA1 leads to the loss of TLE3 binding at these sites in both ENZ 0 and ENZ 3w VCaP cells (Figure 5A). Globally, the loss of FOXA1 decreased TLE3 chromatin binding (Supplementary Figure S11B). At the GRBs, Dex treatment decreased TLE3 chromatin binding at ENZ 3w cells, but to a lesser degree than siFOXA1 (Supplementary Figure S11C). Interestingly, the siFOXA1-UP GRBs showed no change in TLE3 binding after the loss of FOXA1 (Supplementary Figure S11D), suggesting that the increased GR binding is not due to direct chromatin involvement of TLE3. In support of the siFOXA1-induced loss of TLE3 binding at the *NR3C1* promoter, the depletion of either FOXA1 or TLE3 significantly increased the GR transcript levels in VCaP and 22Rv1 cells (Figure 5B-C). A similar increase was observed at the GR protein levels (Supplementary Figure S11E-F). Furthermore, genome-wide gene expression profiling indicated a significant increase in *NR3C1* transcript levels after the depletion of FOXA1 or TLE3 (Supplementary Figure S11G). Moreover, the FOXA1-mediated repression of *NR3C1* was not reciprocal for the regulation of *FOXA1* expression. Even though one GRB is found near the *FOXA1* locus (Supplementary Figure S11H), the depletion of GR with siNR3C1 (Supplementary Figure S3C) did not increase the protein levels of FOXA1 (Supplementary Figure S11I-J). Finally, the increase in GR mRNA levels manifested at the transcriptional regulation, wherein several GR target genes were significantly more Dex-regulated in ENZ 0 and ENZ 3w cells upon the depletion of FOXA1 (Figure 5D-E) or TLE3 (Supplementary Figure 11K-L). A more prominent increase was observed upon depletion of FOXA1 compared to TLE3. Thus, FOXA1 mediates the repression of *NR3C1* expression through corepressor TLE3, restricting the chromatin binding and transcriptional regulatory activity of GR.

**Figure 5.**
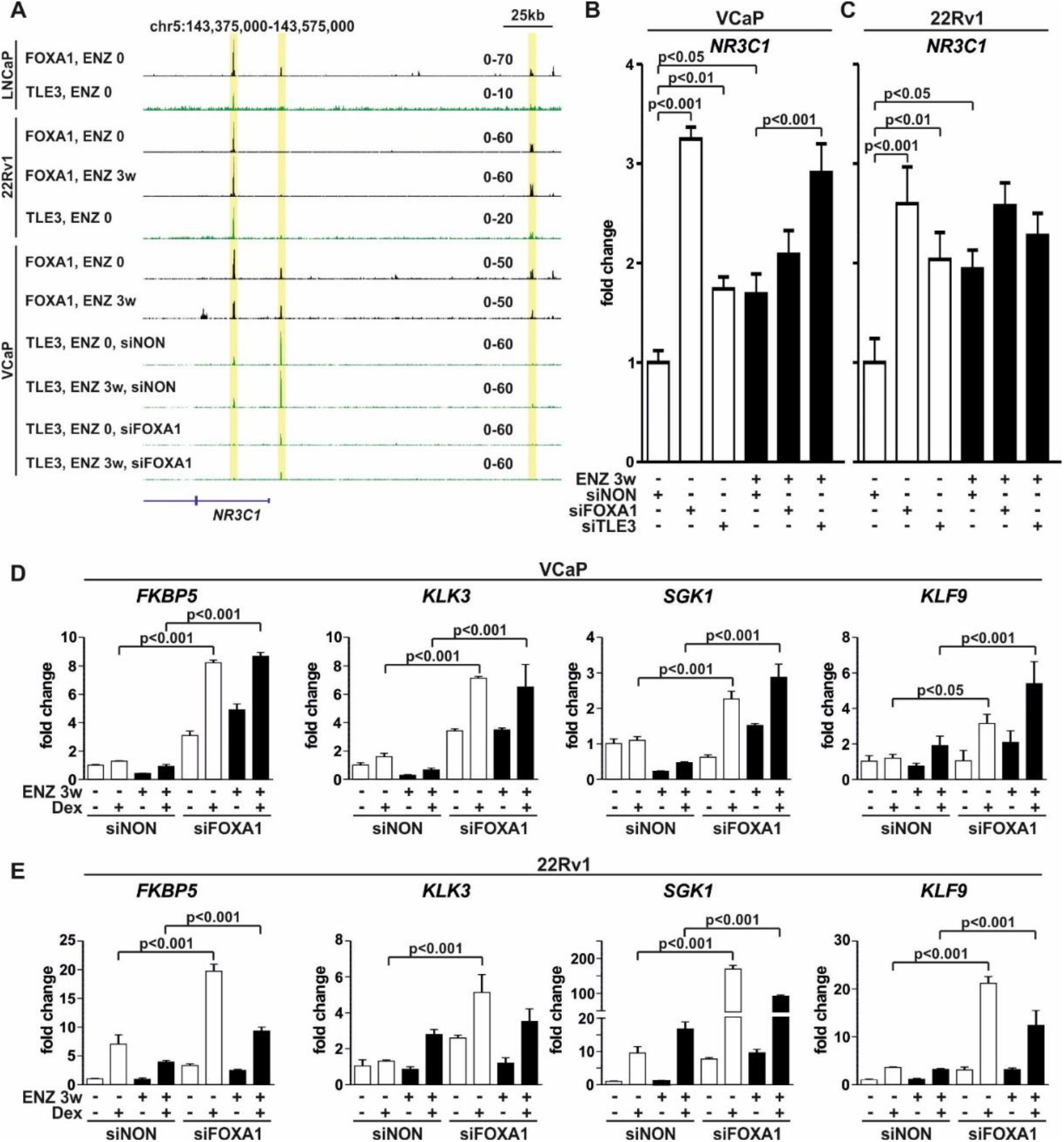
Repression of *NR3C1* expression by FOXA1 is TLE3-mediated. (A) Genome browser tracks of FOXA1 and TLE3 ChIP-seq from LNCaP, 22Rv1, and VCaP cells. FOXA1 ChIP-seq data from 22Rv1 and VCaP ENZ 0 and ENZ 3w cells. TLE3 ChIP-seq data from VCaP ENZ 0 and ENZ 3w cells with or without siFOXA1 treatment. Binding sites highlighted in yellow. All genome browser tracks are normalized to a total of 10 million reads. (B-C) Bar graphs depict *NR3C1* expression levels in VCaP (B) and 22Rv1 (C) ENZ 0 or ENZ 3w cells treated with siNON or siFOXA1 or siTLE3. Bars represent mean ±SD, n=3. Statistical significance calculated using One-way ANOVA with Bonferroni *post hoc* test. (D-E) Bar graphs depict GR target gene analysis of *FKBP5, KLK3, SGK1, KLF9* in VCaP (D) and 22Rv1 (E) ENZ 0 and ENZ 3w cells treated with siNON or siFOXA1 in the presence or absence of Dex. Bars represent mean ±SD, n=3. Statistical significance calculated using One-way ANOVA with Bonferroni *post hoc* test.

### Restriction of *NR3C1* expression by *FOXA1* is not limited to prostate cancer

Previously, it has been shown that, due to the AR-mediated repression of *NR3C1*, the protein levels of GR are higher in AR-negative compared with AR-positive PCa cells (16). To investigate if FOXA1-mediated repression has a potential role in this aspect, we examined *NR3C1* and *FOXA1* expression in recently classified PCa cell types (54). A moderate negative correlation was evident between *NR3C1* and *FOXA1* expression throughout the PCa cell types (Figure 6A). The majority of the CRPC-AR type cells presented high levels of *FOXA1* and low levels of *NR3C1* transcripts, whereas the majority of the CRPC-stem cell like (SCL) type cells showed the opposite. Furthermore, this was not restricted to PCa cell lines, as multiple PCa patient RNA-seq datasets indicated a negative correlation between *NR3C1* and *FOXA1* expression levels (Supplementary Figure S12A). The negative correlation was also observed between *NR3C1* and *TLE3* while *FOXA1* and *TLE3* showed a positive correlation. Interestingly, before receiving ENZ therapy, PCa patients with low levels of *FOXA1* showed significantly higher *NR3C1* transcript levels compared with PCa patients with high levels of *FOXA1* (Figure 6B). Furthermore, ENZ-treated PCa patients displayed similar levels of *NR3C1* as untreated PCa patients with low levels of *FOXA1*. This agrees with our chromatin binding data since GR cistrome is similar between siFOXA1 without ENZ treatment and ENZ 3w cells with intact FOXA1. Moreover, PCa patients with high *NR3C1* and *FOXA1* transcript levels had increased progression-free survival over PCa patients with high *NR3C1* but low *FOXA1* transcript levels (Supplementary Figure S12B-C). This was not seen in PCa patients with low *NR3C1* transcript levels, or if patients were divided based on *TLE3* instead of *FOXA1* expression. These results suggest that GR and FOXA1 have an intertwined relationship in PCa, which impacts the transcriptional regulatory capability of GR.

**Figure 6.**
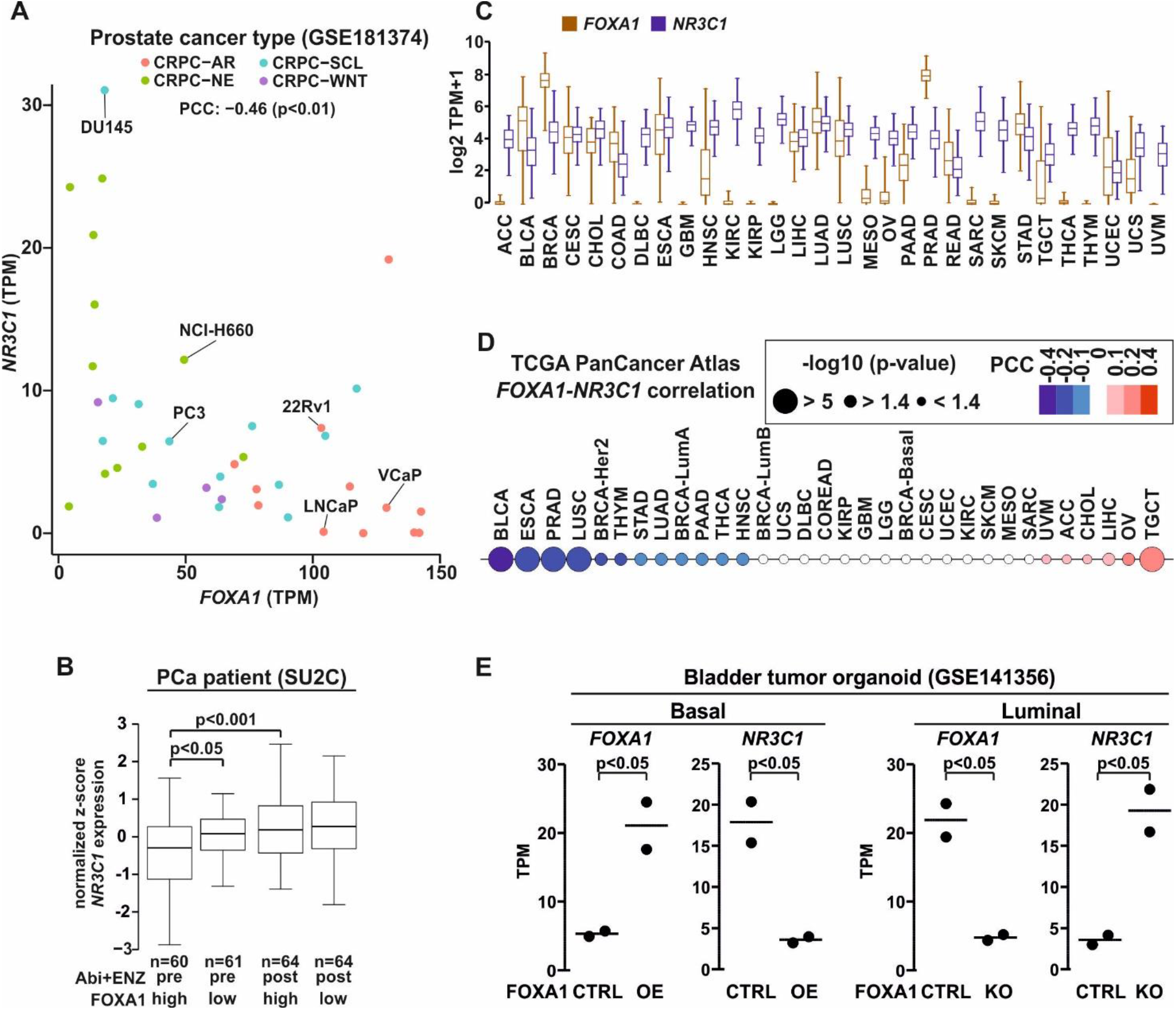
Extensive relationship between the expression of *FOXA1* and *NR3C1* revealed by pan cancer analysis. (A) Scatter plot of *FOXA1* (x-axis) and *NR3C1* (y-axis) expression from RNA-seq in CRPC-AR (red), -NE (green), -SCL (blue), and -WNT (purple) cell types. Data shown as transcripts per million (TPM). Pearson correlation coefficient (PCC) shown above the plot. (B) Box plots of *NR3C1* expression in prostate cancer (PCa) patients divided based on *FOXA1* median expression (high/low), and Abiraterone and ENZ (Abi+ENZ) treatment status (pre/post). Statistical significance calculated using One-way ANOVA with Bonferroni *post hoc* test. (C) Box plots of *FOXA1* (brown) and *NR3C1* (blue) expression from RNA-seq in The Cancer Genome Atlas (TCGA) pan cancer atlas dataset. Data shown as TPM+1. (D) PCC of *FOXA1* and *NR3C1* expression from RNA-seq in TCGA pan cancer atlas dataset. Blue color depicts negative correlation and red color positive correlation. The circle size depicts -log10 p-value. (E) Dot plot represents *FOXA1* and *NR3C1* expression from RNA-seq in bladder tumor basal organoid with overexpression (OE) of FOXA1 or bladder tumor luminal organoid with knockout (KO) of FOXA1. Data shown as TPM. Statistical significance calculated with unpaired t-test.

Since both FOXA1 and GR possess a role beyond PCa (55, 56), we extended the examination of the two TFs to other cancers. *FOXA1* and *NR3C1* are expressed at fluctuating degree between the different tumor types (Figure 6C). TCGA’s Pan Cancer Atlas data indicated that several cancers present negative correlation between *FOXA1* and *NR3C1* transcript levels (Figure 6D). In addition to PCa (PRAD), the most prominent negative correlation was observed in the bladder (BCLA), esophagus (ESCA) and squamous lung (LUSC) cancers, while only a few cancers displayed positive correlation, such as ovarian (OV) and testicular (TGCT) cancers. In general, PRAD, BCLA, ESCA and LUSC showed at least moderate expression of *FOXA1* and *NR3C1* (Figure 6C). Furthermore, CCLE RNA-seq and quantitative proteomics data indicated inverse expression between *FOXA1* and *NR3C1* in several PCa, bladder, esophagus, and squamous lung cancer cell lines (Supplementary Figure S12D). Thus, the FOXA1-mediated repression of GR is likely not limited to PCa. To investigate this more closely, we examined the relationship between FOXA1 and GR in bladder cancer, since it showed the highest negative correlation between the cancers (Figure 6D). For this, we utilized RNA-seq data from bladder tumor organoids (57), with FOXA1 overexpression (OE) in basal and FOXA1 knockout (KO) in luminal tumor organoids. In agreement with our PCa data, FOXA1 OE resulted in a significant decrease while FOXA1 KO manifested in a significant increase of *NR3C1* expression in bladder tumor organoids (Figure 6E). Our results indicate that the *NR3C1* repression by FOXA1 might be a prominent mechanism to restrict the GR action in cancers.

## DISCUSSION

There are several crosstalk mechanisms in cancers through which TFs influence the activity of each other, eventually leading to alterations in gene expression and disease progression (5, 58). Direct TF crosstalk mechanisms are the more studied processes wherein the TFs act at the same chromatin sites, while indirect mechanisms have remained less investigated. However, indirect processes have recently gained more prominence, including such mechanisms as coregulator squelching and TF cascades. In the former, the activation of a TF, such as AR or GR, can sequester coregulators from active enhancers to their binding sites indirectly repressing the pre-existing enhancers (59, 60). In the latter, the activation or repression of initial TF leads to a cascade where the production of another TF is increased, resulting indirectly in the activation of new pathways through the produced TF (61). Intriguingly, the development of antiandrogen resistance through GR in PCa occurs through the TF cascade where the inhibition of AR de-represses *NR3C1* gene expression leading to the GR-mediated replacement of AR signaling (9). Even though the upregulation of GR protein levels is postulated to be a common mechanism of antiandrogen resistance (16), the available unbiased genome-wide information in literature is limited to one atypical cell model. In PCa patients, glucocorticoids are used in the alleviation of therapy-related side effects, and in the compensation of the abiraterone-induced reduction in serum cortisol and to block the compensatory increase in the adrenocorticotropic hormone (5, 9). Thus, it is critical to understand how GR acts in PCa and how it interferes and crosstalks with other signaling pathways. To address this lack of knowledge, we have here utilized two commonly used PCa cell models, VCaP and 22Rv1 cells, and variety of deep sequencing methods to investigate the replacement of AR signaling by GR in the same cellular background.

In both cell lines the GR protein levels, receptor’s chromatin binding and transcriptional activity was significantly increased after treatment with ENZ for 3 weeks. Interestingly, the androgen-independent 22Rv1 cells (41) showed high GR activity in therapy-naïve conditions whereas the androgen-dependent VCaP cells presented a greater increase in GR activity after the ENZ treatment. Thus, as has been previously suggested (16), the loss of androgen dependency increases GR activity in different types of PCa cells. Notably, in VCaP cells the increase in GR chromatin binding after ENZ was distinctly greater than the increase seen in GR protein levels. This variation could be explained *e*.*g*., by the changes in GR post-translational modifications such as phosphorylation (62), or by the increase in the intracellular production of glucocorticoids (17). We observed that the biological pathways regulated by GR in ENZ-treated VCaP and 22Rv1 cells were equivalent to pathways regulated by the AR in therapy-naïve cells. Similar effect has been previously shown on a restricted number of AR-regulated genes (9), however, our results validated this on an unbiased transcriptome-wide level. Moreover, the chromatin replacement of AR by GR was only partial in VCaP and 22Rv1 cells, in agreement with the previously published results in LREX cells (9). Intriguingly, our examination of chromatin landscape pointed out the logic behind the partial replacement. ATAC-seq analyses indicated that the bypass of AR blockade by GR is restricted to pre-accessible chromatin sites. Even though the GR chromatin binding increased in different proportion between VCaP and 22Rv1 cells, the vast majority of GRBs in both cell lines occurred at already open chromatin sites. Moreover, this phenomenon seemed to be rather specific to PCa cells, since the proportion of GRBs at closed *de novo* sites was distinctly higher in other cell types, such as in A549 lung cancer cells and in mouse liver. Interestingly, steroid receptor binding sites that occur at open chromatin sites are postulated to mediate cell type-specific gene regulation (63). This suggested that rather than creating new ones, GR utilizes the cells’ already hard-wired enhancers to mediate antiandrogen resistance. Thus, restricting the activity of these enhancers would inhibit the action of GR.

Previously, the pre-accessibility of GRBs has been associated with the Dex concentration sensitivity (43). However, more recently the enhancer abundance of coactivator p300 prior to GR binding was shown to be better associated with Dex sensitivity than chromatin pre-accessibility (64). Interestingly, ENZ-resistant PCa cells have been shown to be sensitive to p300/CBP inhibition (65). This advocates the importance of p300 at GR-regulated enhancers. Our cistrome and transcriptome analyses showed that the GRBs and GR-regulated target genes in VCaP and 22Rv1 cells were associated not only with p300 but also with other coregulators such as chromatin remodeler SMARCA4 and BET family protein BRD4. Intriguingly, the inhibition of these coregulatory proteins has been shown to restrict AR signaling in PCa cells impeding the disease progression (46, 47, 66, 67). Furthermore, other coregulators such as NCOR2 (68) and CHD1 (69) can contribute to resistance to ADT and AR-targeted therapies. Since many coregulators are shared between AR and GR (3), it is plausible that these coregulators also influence the GR signaling in PCa cells–as our investigations indicated. The inhibition of especially p300 efficiently restricted GR’s transcriptional activity and cell growth of both ENZ-treated and treatment-naïve PCa cells. Thus, our results suggest that the activity of upregulated GR could be restricted *via* p300 inhibition. The influence of p300 inhibition on the genome-wide glucocorticoid signaling and PCa cells’ hard-wired enhancers is currently underway, and it will reveal the extent of inhibitor utility in restricting GR-mediated antiandrogen resistance.

FOXA1 is a well-known and important regulator of AR signaling in PCa cells (50, 70), and it also modulates the action of GR (51). The occurrence of FOXA1 motif and FOXA1 chromatin binding was clearly detectable at the GRBs in both VCaP and 22Rv1 cells. Furthermore, GRBs in LREX cells display the enrichment of FOXA1 motif (9), and on a single-cell level the ENZ-treatment did not change accessibility at sites with FOXA1 motif in LNCaP and its derivative cells (71). Since GRBs are found at these types of open chromatin sites, FOXA1 is implied to contain a crucial role on GR signaling in PCa cells. Surprisingly, our analyses indicated that the depletion of FOXA1 potentiated GR signaling, and the potentiation was due to FOXA1-mediated repression of *NR3C1* expression. It has been shown that FOXA1 represses *AR* expression (50). However, in the case of GR, the previous study did not observe the repression of *NR3C1* or the potentiation of GR signaling after FOXA1 depletion, but rather modulation of receptor binding (51). This difference is most likely due to the utilized LNCaP-1F5 cells. LNCaP cells do not naturally express GR until the cells are exposed to ENZ (16, 42), whereas LNCaP-1F5 cells express stably integrated rat GR driven by the CMV promoter (72). Thus, the natural cellular milieu is required to observe the FOXA1-mediated repression of *NR3C1* expression. What remains unclear is the role of FOXA1 occupancy at the GRBs. The slight modulation of GRBs that we observed after the depletion of FOXA1 was lower than what was previously shown (51). This can be attributed to the utilized cell models as discussed above. The study postulated that the large number of FOXA1-independent receptor binding sites may collaborate with other TFs. Interestingly, our analyses indicated that HOXB13 and GATA-family motifs were enriched at GRBs, although to a lesser degree than FOXA1 motifs. HOXB13 has not been implicated in GR signaling, but GATA2 can influence *NR3C1* expression in ENZ-treated PCa cells (73). However, the role of GATA2 at GRBs has not been shown. Nonetheless, AR signaling is modulated by GATA2 (74), implying its potential involvement in the regulation of GR’s transcriptional activity in PCa. It has also been suggested that FOXA1 shifts AR equilibrium from high-to low-affinity binding sites, *i*.*e*., from sites with canonical AREs to sites with FOXA1 motifs (50). Our analyses show that GRBs that appear only after ENZ treatment and FOXA1 depletion, contain high frequency of GREs, representing high-affinity sites. Thus, FOXA1 could shift the binding equilibrium of GR but it is masked by the more global potentiation of GR activity.

Although GR activity is similarly induced by ENZ treatment or by the depletion of FOXA1, the mechanistic repression of *NR3C1* occurs in a dissimilar way. While *NR3C1* repression *via* AR is postulated to function through EZH2 (15), we discovered that corepressor TLE3 is crucially involved in FOXA1-mediated repression. Previous investigation has indicated that TLE3 is recruited to chromatin by FOXA1, and the corepressor is more efficient in creating closed chromatin when it is recruited by a TF (52). Furthermore, TLE3 has been shown to mediate the repression of *NR3C1* in PCa cells (53). Our investigations indicate that FOXA1 is the main recruiter of TLE3 to *NR3C1* locus, wherein TLE3 chromatin binding is lost after the depletion of FOXA1. Furthermore, FOXA1 has a wide impact on TLE3, since its protein levels and genome-wide chromatin occupancy are decreased upon FOXA1 depletion. Beyond GR and FOXA1 in PCa, TLE3 has been shown to restrict estrogen receptor signaling (75), and FOXC1-mediated recruitment of TLE3 limits monocyte enhancer activity (76). This suggests an extensive role of TLE3 in restricting steroid receptor signaling, whereas FOX-family members could be involved in the mediation of TLE3’s chromatin activity. Interestingly, the FOXA1-mediated repression of *NR3C1 i*.*e*., the inverse correlation between the two TFs, is not restricted to PCa but can be observed in other cancers. This is especially prevalent in bladder cancer, where our reanalysis of published data (57) indeed indicated the repression of *NR3C1* expression by FOXA1. Both FOXA1 and GR have been implicated in bladder cancer (56, 57). This demonstrates that the *NR3C1* repression *via* FOXA1 could be a widespread mechanism to restrict GR activity in cancers. In support of the broad repressive role of FOXA1, interferon signaling is suppressed in both PCa and bladder cancers by FOXA1 (55).

While the GR acts as an oncogene mediating antiandrogen resistance, the receptor has originally been postulated to function in a tumor suppressive manner (77). Thus, GR can act either as a tumor suppressor or as an oncogene, likely depending on the cellular context. This duality has been recognized in several cancers (56). Intriguingly, PCa cells display a distinct expression of GR (16, 54), with the highest expression at the AR-negative stem cell-like and neuroendocrine subtypes. In the case for the latter subtype, GR has been implicated in neuroendocrine PCa through the regulation of MYCN (78). Thus, GR likely has an extensive role in AR-negative PCa subtypes which warrants further investigation, especially since these subtypes represent around half of the PCa patients (54). Taken together, our research presented here demonstrates that *NR3C1* expression is repressed by the AR and FOXA1, and inhibition of either one leads to the upregulated GR and replacement of AR occupancy at pre-accessible chromatin sites (Figure 7). The importance of open chromatin and specific coregulators on the GR action in PCa points out new avenues to oppose steroid receptor-mediated drug resistance.

**Figure 7.**
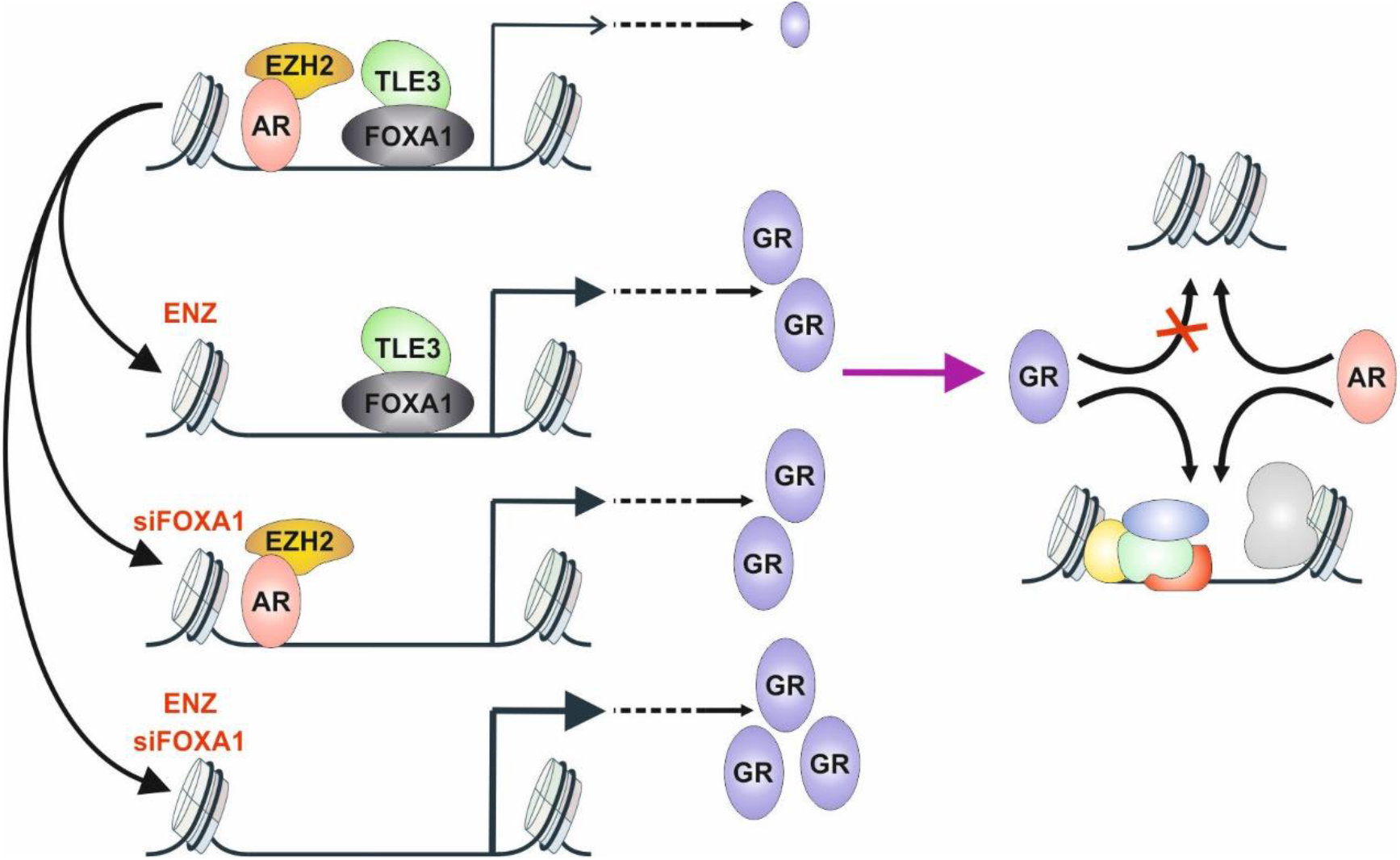
Model of AR and FOXA1 mediated repression of *NR3C1* expression. The expression of *NR3C1* is repressed by AR through EZH2, and by FOXA1 through TLE3. Inhibition of AR by ENZ and/or depletion of FOXA1 by siRNA (siFOXA1) leads to de-repression of *NR3C1*. De-repression of *NR3C1* leads to increased GR protein levels, and augmentation of GR transcriptional activity. GR occupies AR binding sites at pre-accessible chromatin sites enriched with coregulator proteins.

## DATA AVAILABILITY

The generated ChIP-seq, RNA-seq and ATAC-seq datasets have been submitted to the NCBI Gene Expression Omnibus database (http://www.ncbi.nlm.nih.gov/geo/). Accession code: GSE214757.

## FUNDING

This work was supported by the Academy of Finland; Cancer Foundation Finland; Sigrid Jusélius Foundation.

## ACKNOWLEDGEMENTS

We thank Dr Leena Latonen for providing the 22Rv1 cells, Dr Janne Capra for help with image analysis, and Merja Räsänen, Eija Korhonen and Saara Pirinen for technical assistance. The EMBL GeneCore team (Heidelberg, Germany) is greatly acknowledged for deep sequencing services. This work was carried out with the support of UEF Cell and Tissue Imaging Unit, University of Eastern Finland, Finland, and Biocenter Finland. The computational analyses were performed on servers provided by UEF Bioinformatics Center, University of Eastern Finland, Finland.

## AUTHOR CONTRIBUTIONS

Data acquisition and analysis: LH, JH, NA, EAN, VP. Manuscript writing: LH, VP. Research funding and supervision: JJP, VP. All authors read and approved the manuscript.

## CONFLICTS OF INTEREST

None declared.

## Notes

### Competing Interest Statement

The authors have declared no competing interest.

