## Supplemental material for "Chromatin Accessibility and Pioneer Factor FOXA1 Shape Glucocorticoid Receptor Action in Prostate Cancer"

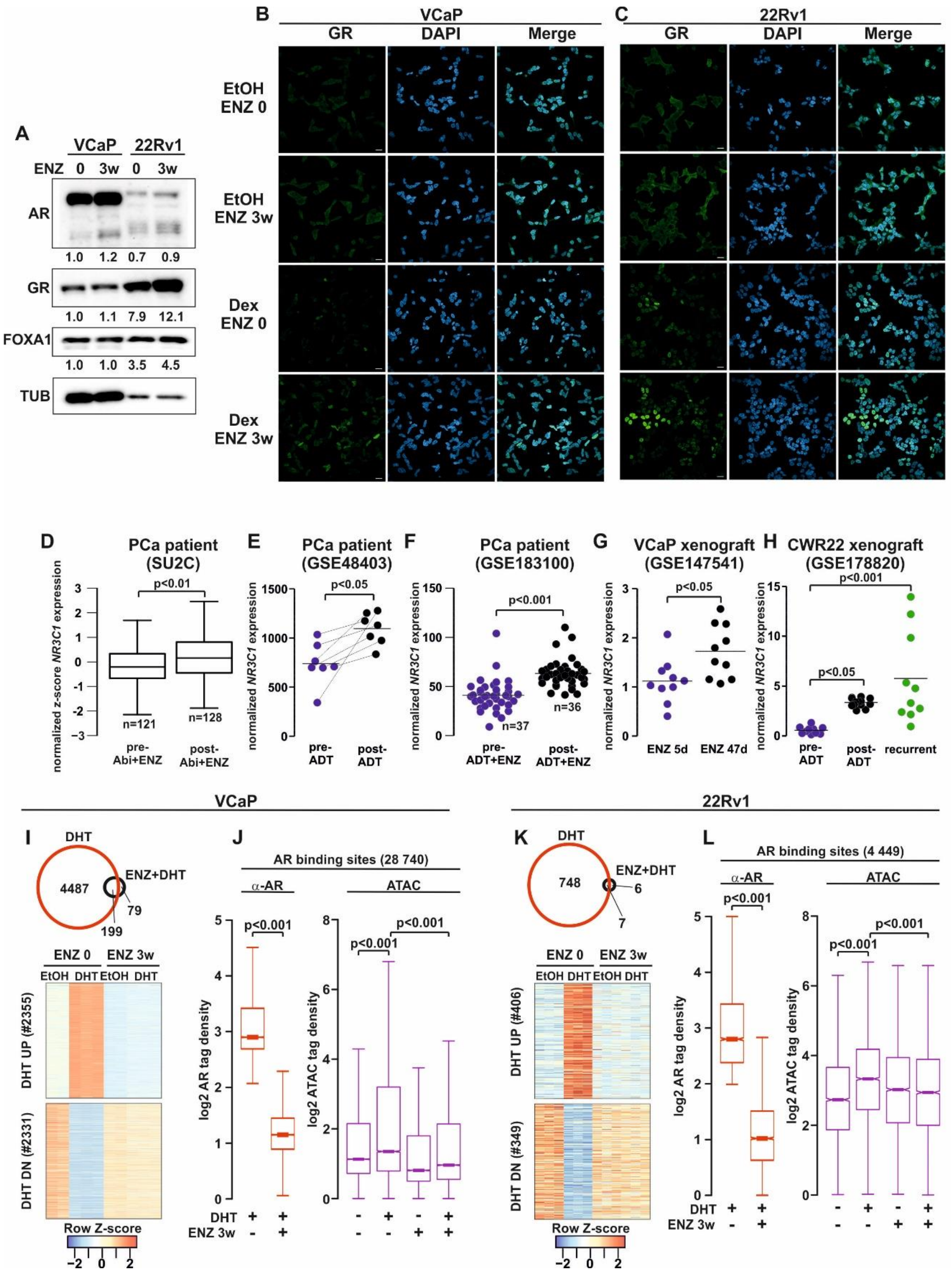


**Supplementary Figure S1. Increase of GR levels and repression of AR signaling by enzalutamide.** (A) Immunoblotting of AR, GR, FOXA1, and tubulin (TUB) protein levels in VCaP and 22Rv1 ENZ 0 and ENZ 3w cells. Quantification of AR, GR, and FOXA1 levels to TUB is shown below the immunoblot. (B-C) GR immunofluorescence (IF) images from VCaP (B) and 22Rv1 (C) cells. The cells were treated with vehicle (EtOH), ENZ 3w, Dex, or ENZ 3w and Dex. DAPI was used as a marker for nuclei. Scale bar, 20 µm. (D-H) Transcript levels of *NR3C1* by RNA-seq from (D) PCa patients pre- or post-exposure with abiraterone (Abi) and/or enzalutamide (ENZ), (E) matched PCa patients pre- or post-exposure with androgen deprivation therapy (ADT), (F) PCa patients pre- or post-exposure with ADT and ENZ, (G) VCaP xenograft after 5 (5d) or 47 days (47d) exposure to ENZ, and (H) CWR22 xenograft pre- or post-exposure with ADT and recurrent samples. Data accession number is indicated for each dataset. (I-L) Inhibition of AR signaling by ENZ 3w in (I-J) VCaP and 22Rv1 (K-L) cells at the level of (I, K) AR target genes regulation by RNA-seq, and (J, L) AR chromatin binding by ChIP-seq and AR-induced chromatin accessibility by ATAC-seq. (I, K) Venn diagrams depict the overlap and number of regulated target genes, and heatmaps are displayed as row Z-score. (J, L) Box plots represent the normalized log2 tag density of AR ChIP-seq and ATAC-seq at AR binding sites. The number of AR binding sites is indicated for each cell lines. All box plots are normalized to a total of 10 million reads. Statistical significance calculated with unpaired t-test (2 conditions) or using One-way ANOVA with Bonferroni *post hoc* test (3 conditions).

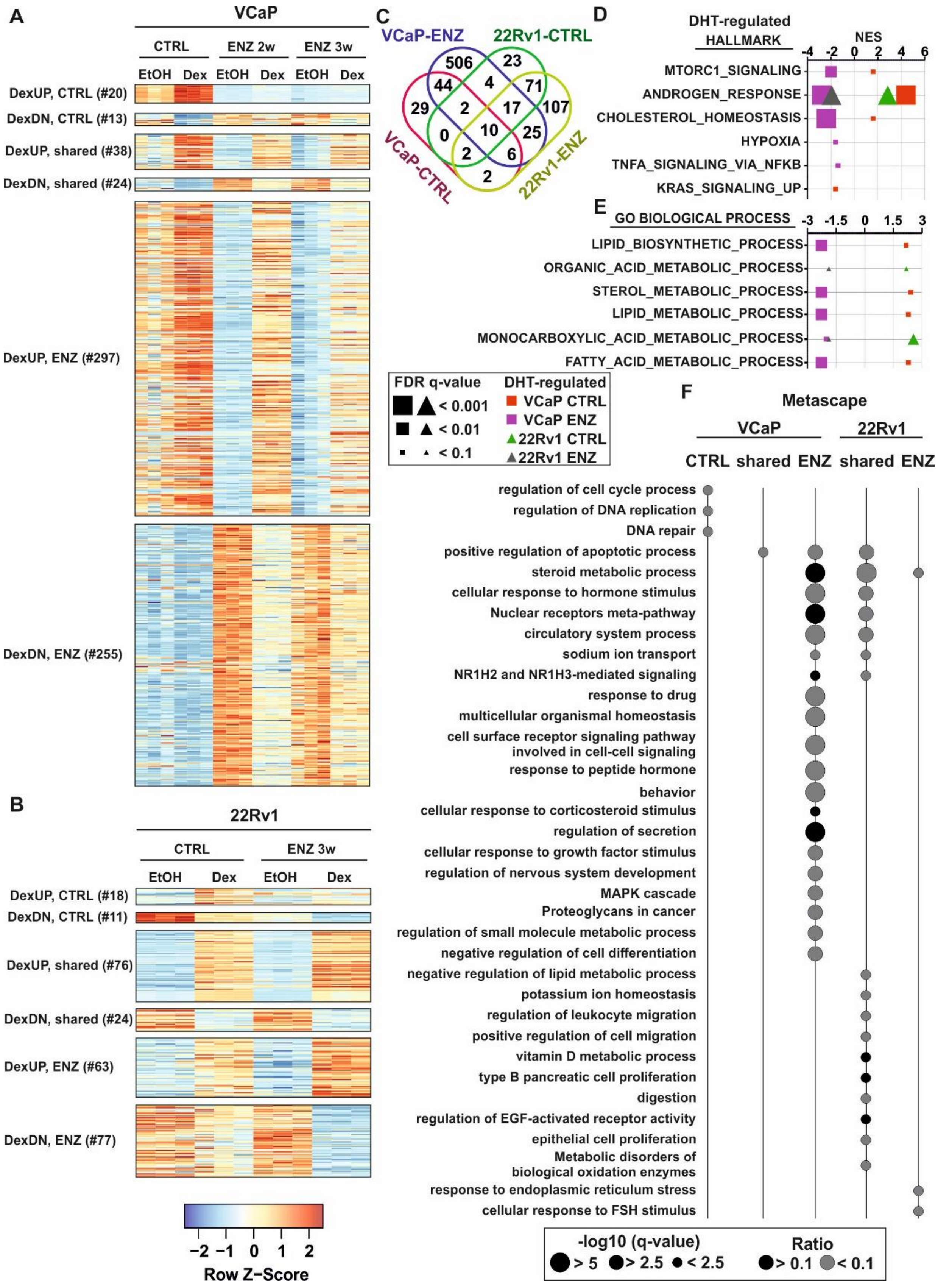


**Supplementary Figure S2. RNA-seq of Dex-regulated target genes in VCaP and 22Rv1 cells.** (A-B) Heatmaps of Dex-regulated genes in (A) VCaP and (B) 22Rv1 cells before and after indicated ENZ treatment. Gene clusters separated based on target gene regulation only at before ENZ (CTRL), in both before and after ENZ (shared), and only after ENZ treatment. (C) Venn diagrams overlap of Dex-regulated genes between VCaP and 22Rv1 cells before and after ENZ treatment. (D-E) Gene set enrichment analysis (GSEA) of DHT-regulated genes from VCaP or 22Rv1 before (VCaP, red square; 22Rv1, green triangle) or after (VCaP, purple square; 22Rv1, grey triangle) ENZ treatment at hallmark (D) or GO biological process (E) pathways. Data shown as normalized enrichment score (NES) with square and triangle size depicting FDR q-value. (F) Metascape pathway analysis of Dex-regulated genes from VCaP and 22Rv1 cells from gene clusters. Circle size depicts -log10 q-value, and black circle indicated over 0.1 and grey circle less than 0.1 gene ratio.

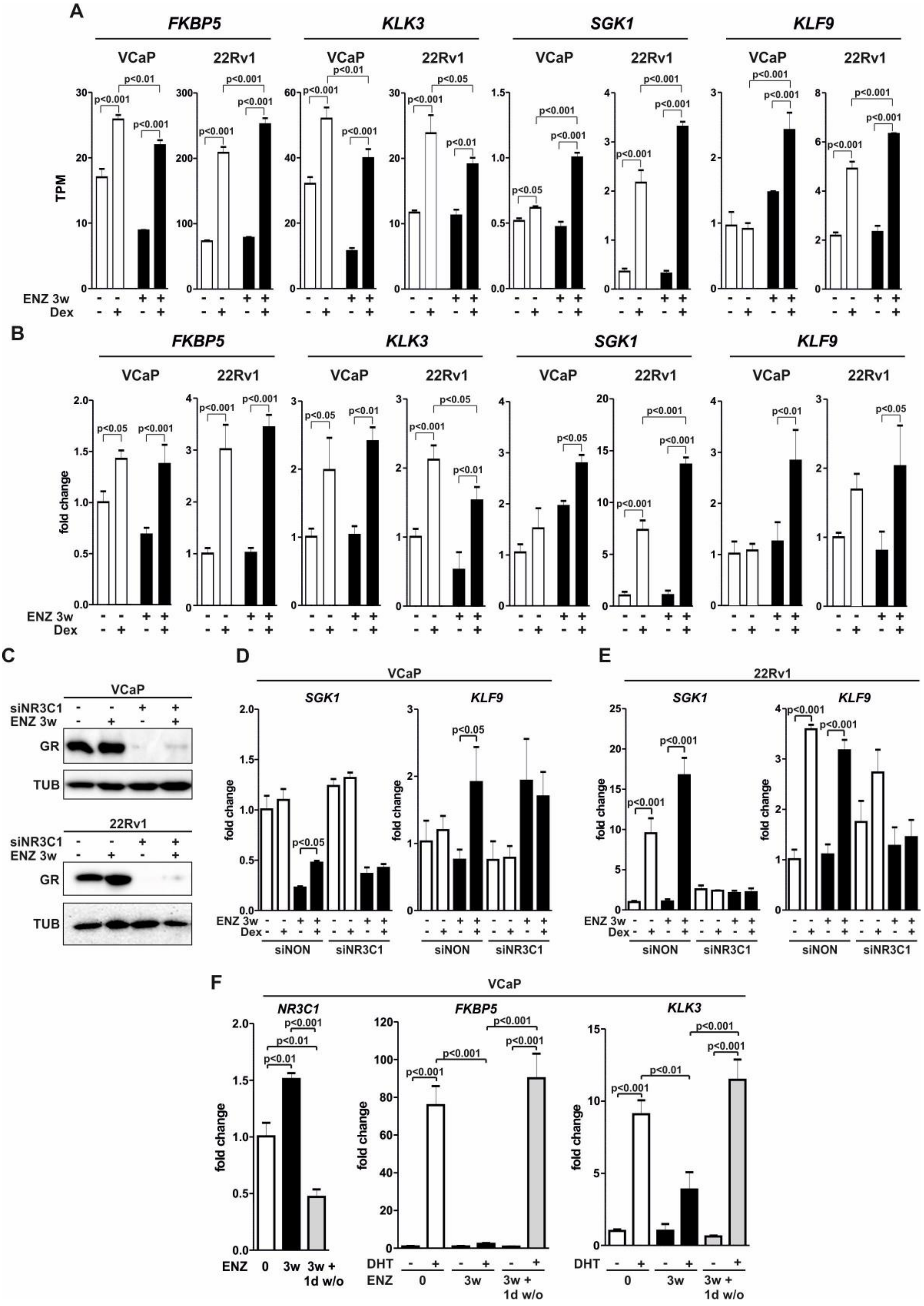


**Supplementary Figure S3. Validation of Dex-regulated genes in enzalutamide treated VCaP and 22Rv1 cells.** (A) Bar graphs depict transcript per million (TPM) of *FKBP5*, *KLK3*, *SGK1*, *KLF9* in VCaP and 22Rv1 ENZ 0 and ENZ 3w cells in the presence or absence of Dex from RNA-seq data. (B) Bar graphs depict GR target gene analysis of *FKBP5*, *KLK3*, *SGK1*, *KLF9* in VCaP and 22Rv1 ENZ 0 and ENZ 3w cells in the presence or absence of Dex. (C) Immunoblotting of GR and TUB protein levels in VCaP (upper) and 22Rv1 (lower) ENZ 0 and ENZ 3w cells treated with siNON or siNR3C1. (D-E) Bar graphs depict GR target gene analysis of *SGK1* and *KLF9* in VCaP (D) and 22Rv1 (E) ENZ 0 and ENZ 3w cells treated with siNON or siNR3C1 in the presence or absence of Dex. (F) Bar graphs depict target gene analysis of NR3C1, *FKBP5* and *KLK3* in VCaP ENZ 0 cells, ENZ 3w cells, or cells treated with ENZ 3w followed by 1 day without ENZ (1d w/o) in the presence or absence of DHT. Bars represent mean ±SD, n=3. Statistical significance calculated using One-way ANOVA with Bonferroni *post hoc* test.

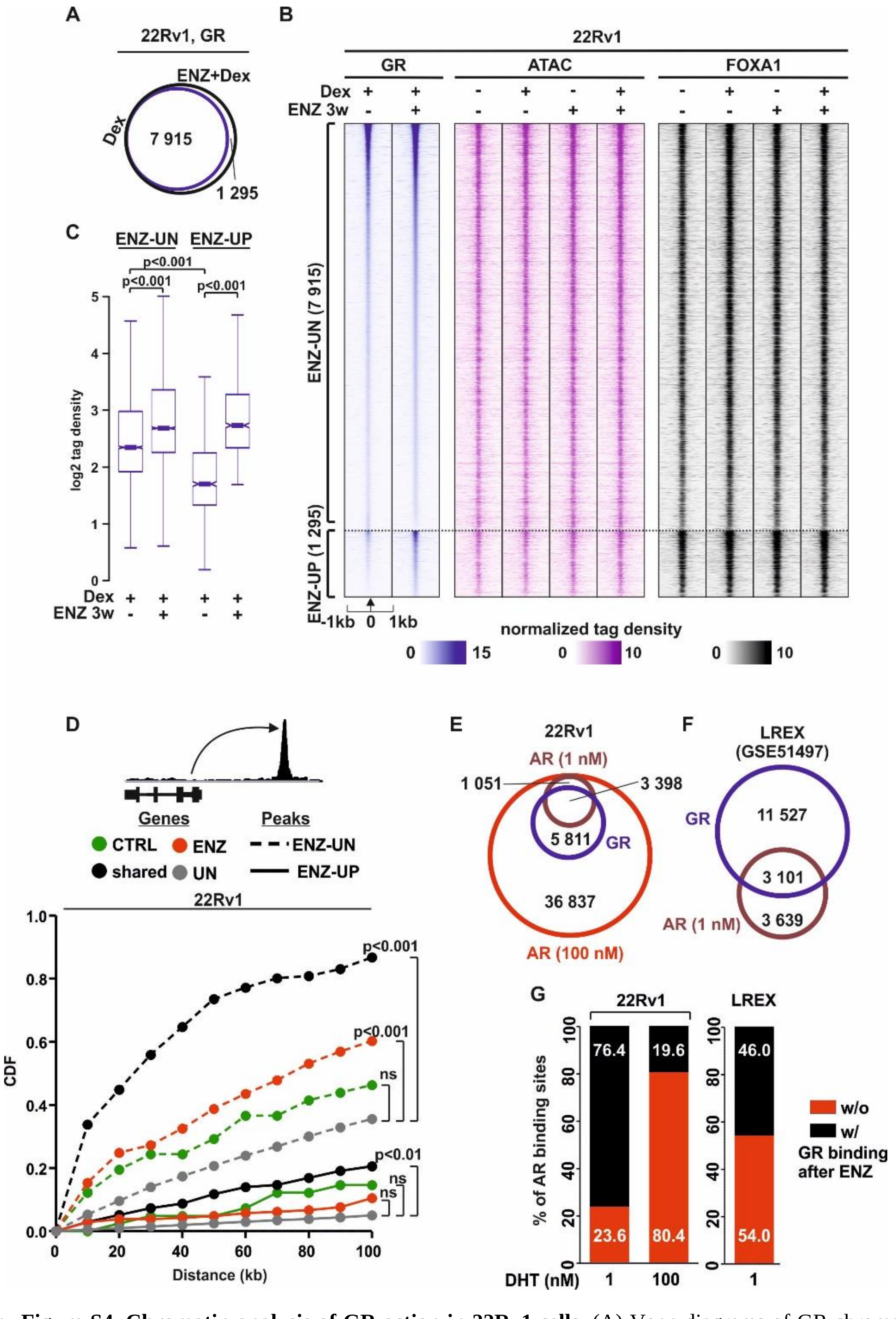


**Supplementary Figure S4. Chromatin analysis of GR action in 22Rv1 cells.** (A) Venn diagrams of GR chromatin binding in 22Rv1 cells from GR ChIP-seq after Dex (blue circle) or ENZ+Dex (black circle) treatment. (B) GR ChIP-seq, ATAC-seq, and FOXA1 ChIP-seq profiles at ENZ-UN and ENZ-UP sites in 22Rv1 ENZ 0 and ENZ 3w cells. ENZ-UN represents unchanged and ENZ-UP increased GR binding sites. Each heatmap represents ±1 kb around the center of the GR peak. Binding intensity (tags per bp per site) scale is noted below on a linear scale. (C) Box plots represent the normalized log2 tag density of GR ChIP-seq at ENZ-UN and ENZ-UP sites. Statistical significance calculated using One-way ANOVA with Bonferroni *post hoc* test. (D) Association of Dex-regulated genes in 22Rv1 ENZ 0 and ENZ 3w cells to ENZ-UN and ENZ-UP GR binding sites. Different subsets of Dex-regulated genes are color coded, and ENZ-UN depicted as dotted line and ENZ-UP as solid line. Statistical significance calculated with Kolmogorov–Smirnov test. (E) Venn diagrams of GR and AR chromatin binding in 22Rv1 cells from GR ChIP-seq (blue circle) and AR ChIP-seq treated with 1 nM (magenta circle) or 100 nM (red circle) of DHT. (F) Venn diagrams of GR and AR chromatin binding in LREX cells from GR ChIP-seq (blue circle) and AR ChIP-seq treated with 1 nM (magenta circle) of DHT. (G) Bar graphs depict percentage overlap of AR binding sites with (black) or without (red) GR binding in 22Rv1 and LREX cells. All heatmaps and box plots are normalized to a total of 10 million reads.

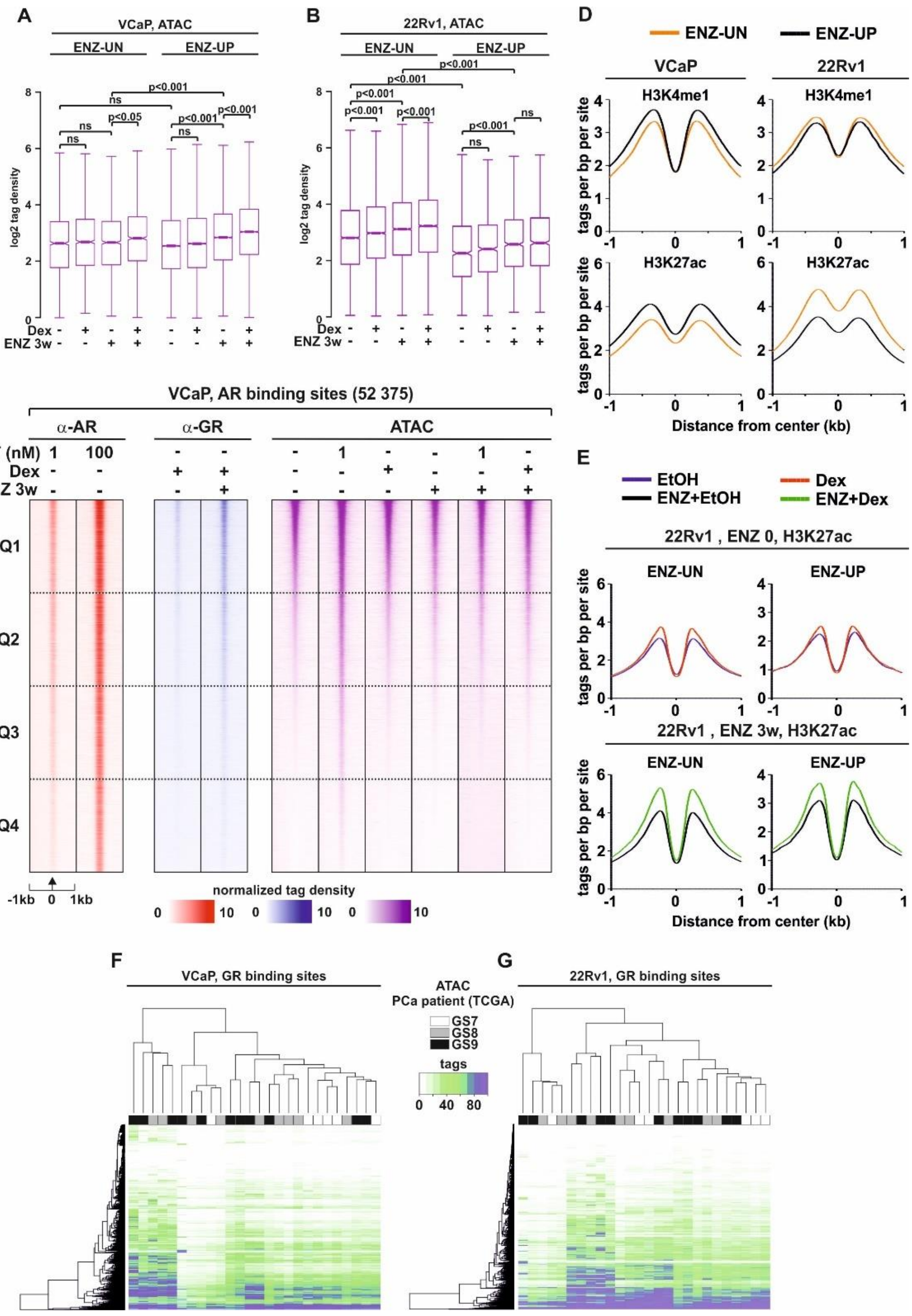


**Supplementary Figure S5. Chromatin accessibility and histone modification enrichment at GR binding sites.** (A-B) Box plots represent the normalized log2 tag density of ATAC-seq at ENZ-UN and ENZ-UP sites in (A) VCaP and (B) 22Rv1 cells. Statistical significance calculated using One-way ANOVA with Bonferroni *post hoc* test. (C) AR ChIP-seq, GR ChIP-seq and ATAC-seq profiles at AR binding sites in VCaP cells. AR binding sites were sorted based on pre-accessibility to quartiles with Q1 the most and Q4 the least pre-accessible. Each heatmap represents ±1 kb around the center of the AR peak. Binding intensity (tags per bp per site) scale is noted below on a linear scale. (D) Aggregate plots represent the binding intensity (tags per bp per site) of H3K4me1 (upper) and H3K27ac (lower) ChIP-seq at ENZ-UN and ENZ-UP sites in untreated VCaP and 22Rv1 cells. (E) Aggregate plots represent the binding intensity (tags per bp per site) of H3K27ac ChIP-seq at ENZ-UN and ENZ-UP sites in 22Rv1 ENZ 0 (upper) and ENZ 3w (lower) cells. Each aggregate plot represents ±1 kb around the center of the GR peak. (F-G) Heatmap of ATAC-seq from PCa patients at GR binding sites in (F) VCaP and (G) 22Rv1 cells. Columns represent different patients with color code depicting patient Gleason score (GS). Rows represent individual GR binding sites. Binding intensity scale is shown on a linear scale. All heatmaps, aggregate and box plots are normalized to a total of 10 million reads.

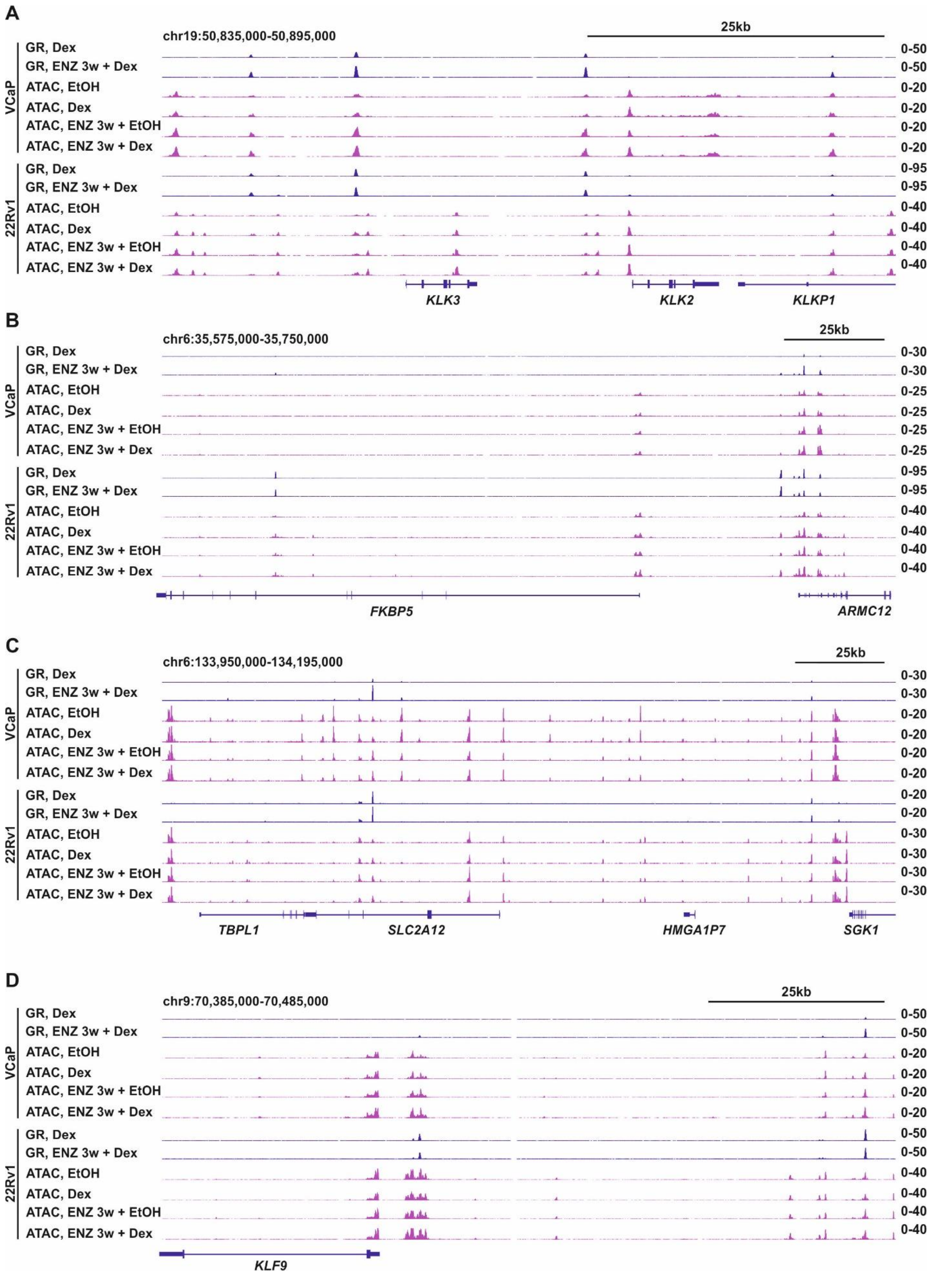


**Supplementary Figure S6. Example genome browser tracks.** Genome browser tracks of (A) *KLK3*, (B) *FKBP5*, (C) *SGK1*, and (D) *KLF9* loci depicting GR ChIP-seq and ATAC-seq in VCaP and 22Rv1 ENZ 0 and ENZ 3w cells treated with or without Dex. All genome browser tracks are normalized to a total of 10 million reads.

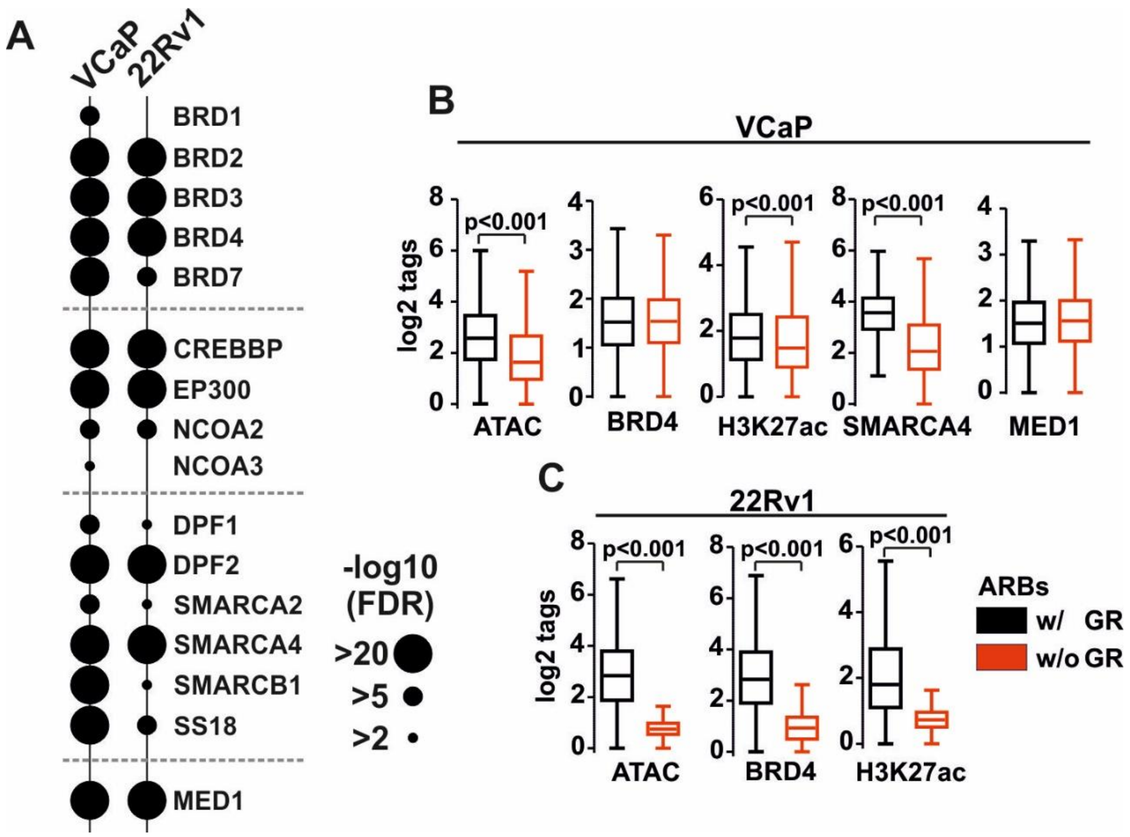


**Supplementary Figure S7. Enrichment of coregulators at AR binding sites with or without GR occupancy.** (A) TcoF gene set enrichment of Dex-regulated genes in VCaP and 22Rv1 cells. Circle size depicts -log10 FDR value. (B) Box plots represent the normalized log2 tag density of ATAC-seq, BRD4 ChIP-seq, H3K27ac ChIP-seq, SMARCA4 ChIP-seq, and MED1 ChIP-seq at AR binding sites with (w/; black) or without (w/o; red) GR occupancy in VCaP cells. (C) Box plots represent the normalized log2 tag density of ATAC-seq, BRD4 ChIP-seq, and H3K27ac ChIP-seq at AR binding sites with (w/; black) or without (w/o; red) GR occupancy in 22Rv1 cells. All box plots are normalized to a total of 10 million reads. Statistical significance calculated with unpaired t-test.

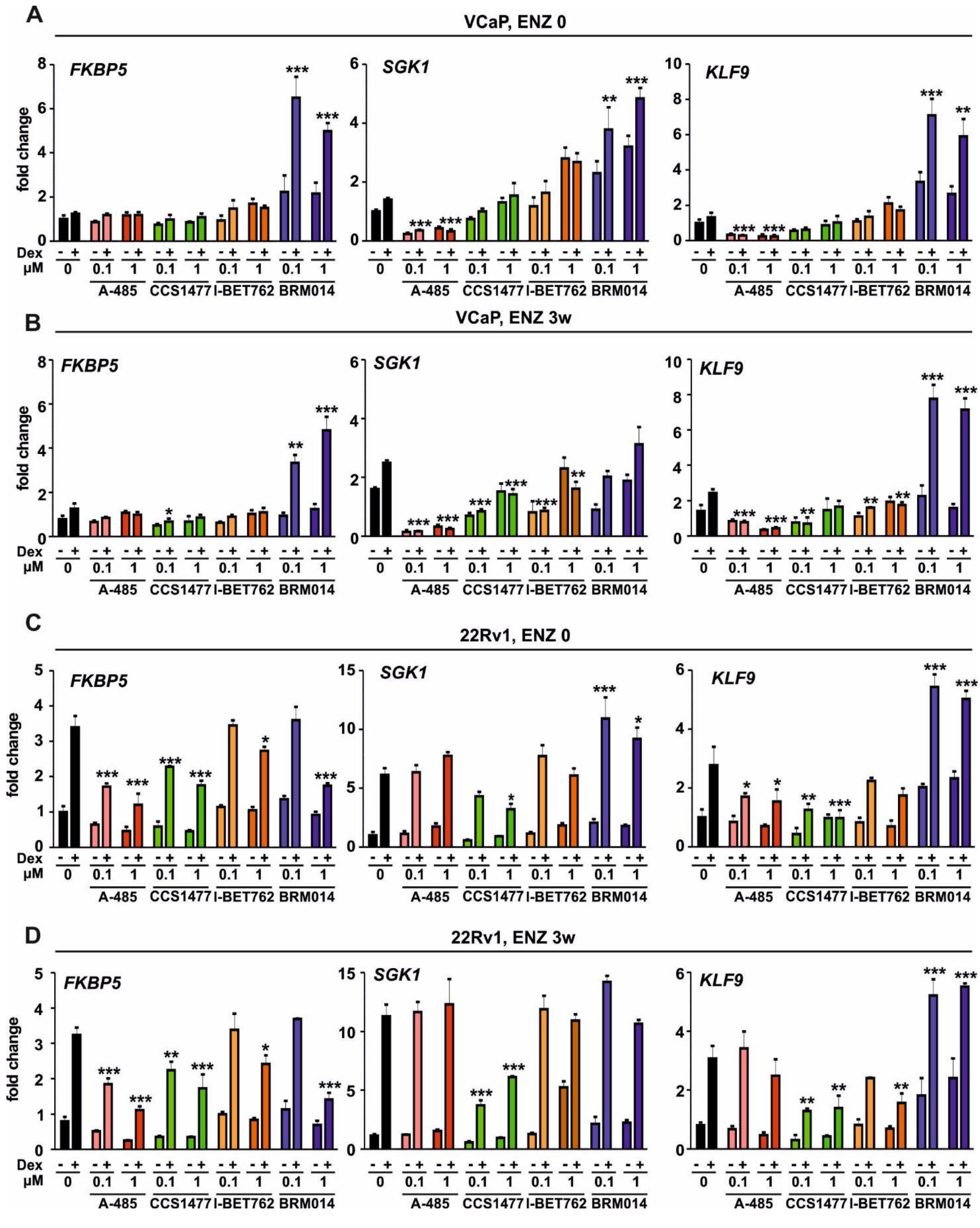


**Supplementary Figure S8. Influence of coregulator inhibitors on GR target gene expression.** (A-D) Bar graphs depict *FKBP5*, *SGK1*, and *KLF9* gene expression analysis in VCaP ENZ 0 (A), VCaP ENZ 3w (B), 22Rv1 ENZ 0 (C), and 22Rv1 ENZ 3w (D) cells treated with or without Dex in the presence or absence of 0.1 or 1 µM of indicated inhibitor. Bars represent mean ±SD, n=3. Statistical significance calculated using One-way ANOVA with Bonferroni *post hoc* test. *, $p<0.05$; **. $p<0.01$; ***, $p<0.001$.

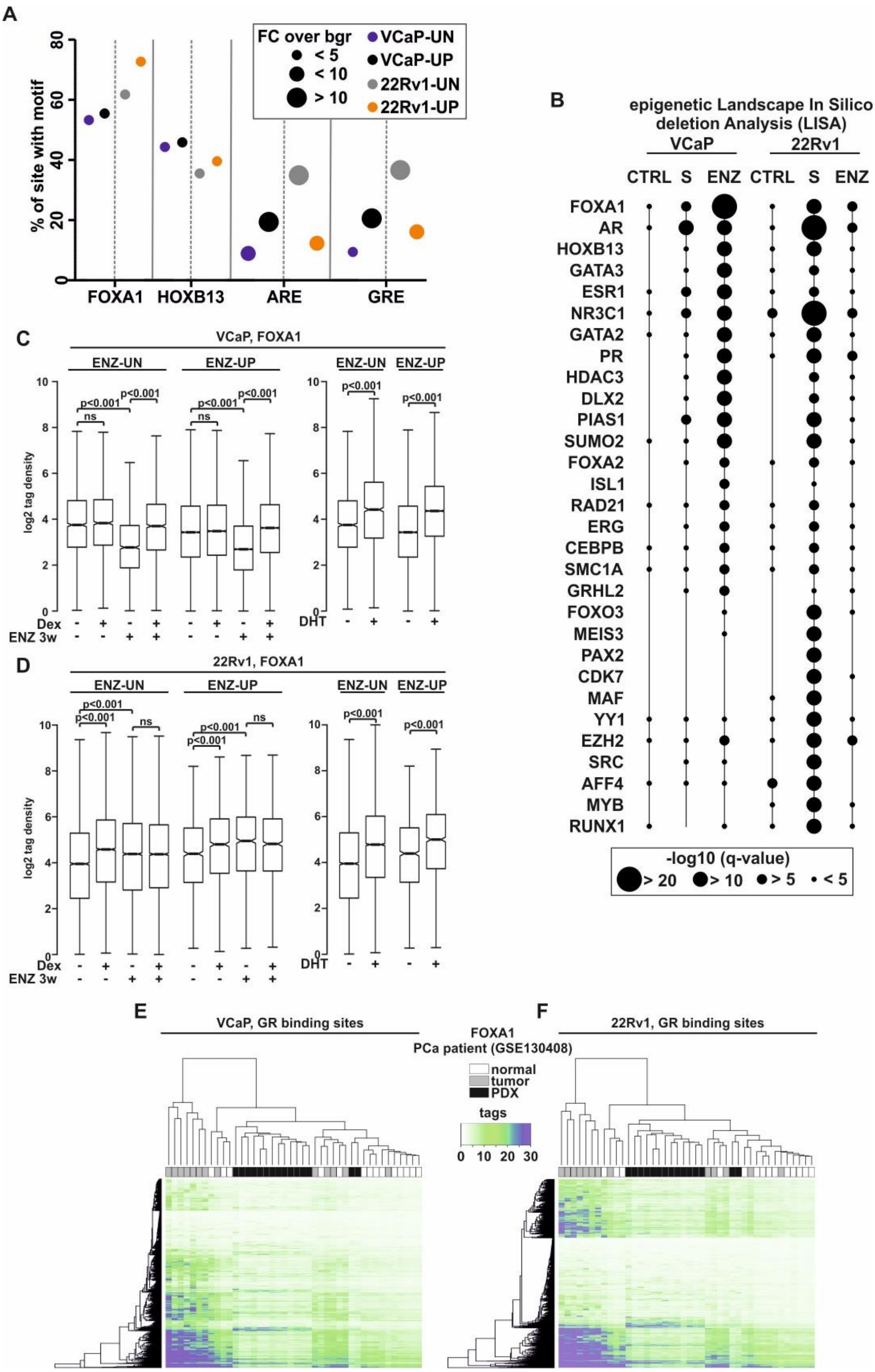


**Supplementary Figure S9. Analysis of FOXA1 enrichment at GR binding sites.** (A) *De novo* motif enrichment of FOXA1, HOXB13, ARE, and GRE motifs at the ENZ-UN (VCaP, blue; 22Rv1 grey) and ENZ-UP (VCaP, black; 22Rv1, orange) sites in VCaP and 22Rv1 cells. Y-axis represents percentage of sites with motifs, and circle size depicts fold change (FC) over the background. (B) Epigenetic landscape in silico deletion analysis (LISA) of Dex-regulated genes from VCaP and 22Rv1 cells from indicated gene clusters. Circle size depicts -log10 q-value. (C-D) Box plots represent the normalized log2 tag density of FOXA1 ChIP-seq at ENZ-UN and ENZ-UP sites in (C) VCaP and (D) 22Rv1 cells. Statistical significance calculated using One-way ANOVA with Bonferroni *post hoc* test. (E-F) Heatmap of FOXA1 ChIP-seq from PCa patients at GR binding sites in (E) VCaP and (F) 22Rv1 cells. Columns represent different patients with color code depicting sample type. Rows represent individual GR binding sites. Binding intensity scale is shown on a linear scale. All heatmaps and box plots are normalized to a total of 10 million reads.

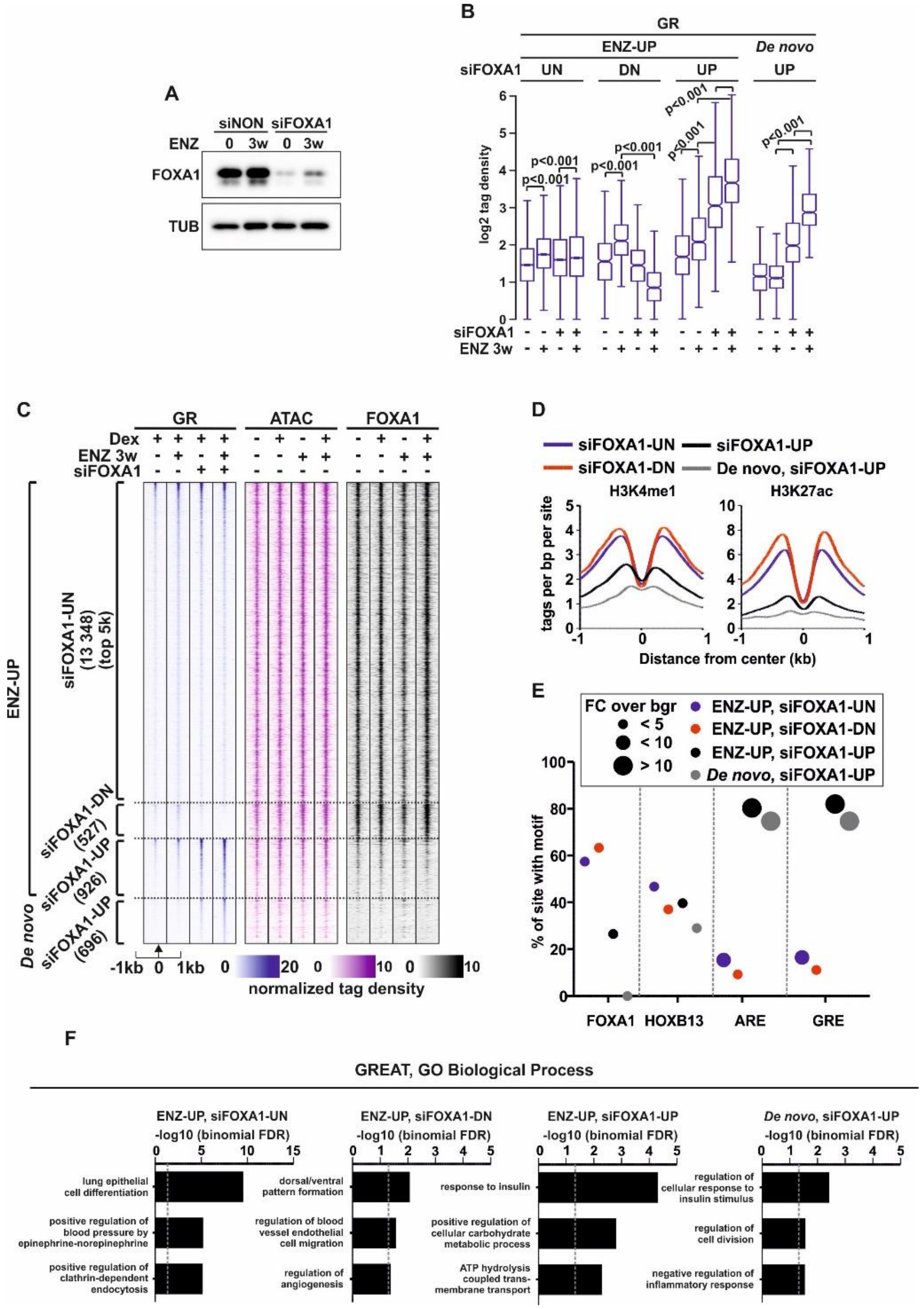


**Supplementary Figure S10. Depletion of FOXA1 increases GR chromatin binding.** (A) Immunoblotting of FOXA1 and TUB protein levels in VCaP ENZ 0 and ENZ 3w cells treated with siNON or siFOXA1. (B) Box plots represent the normalized log2 tag density of GR ChIP-seq at ENZ-UP siFOXA1-UN, ENZ-UP siFOXA1-DN, ENZ-UP siFOXA1-UP, and *de novo* siFOXA1-UP sites in VCaP cells. Statistical significance calculated using One-way ANOVA with Bonferroni *post hoc* test. (C) GR ChIP-seq, ATAC-seq, and FOXA1 ChIP-seq profiles at ENZ-UP siFOXA1-UN, ENZ-UP siFOXA1-DN, ENZ-UP siFOXA1-UP, and *de novo* siFOXA1-UP sites in VCaP cells. Each heatmap represents ±1 kb around the center of the GR peak. Binding intensity (tags per bp per site) scale is noted below on a linear scale. (D) Aggregate plots represent the binding intensity (tags per bp per site) of H3K4me1 (left) and H3K27ac (right) ChIP-seq at ENZ-UP siFOXA1-UN, ENZ-UP siFOXA1-DN, ENZ-UP siFOXA1-UP, and *de novo* siFOXA1-UP sites in VCaP cells. (E) *De novo* motif enrichment of FOXA1, HOXB13, ARE, and GRE motifs at the ENZ-UP siFOXA1-UN (blue), ENZ-UP siFOXA1-DN (red), ENZ-UP siFOXA1-UP (black), and *de novo* siFOXA1-UP (grey) sites in VCaP cells. Y-axis represents percentage of sites with motifs, and circle size depicts fold change (FC) over the background. (F) GREAT enrichment analysis at GO Biological Process pathways for of ENZ-UP siFOXA1-UN, ENZ-UP siFOXA1-DN, ENZ-UP siFOXA1-UP, and *de novo* siFOXA1-UP sites in VCaP cells. Each bar depicts -log10 binomial FDR values, and grey dashed line depicts binomial FDR value of 0.05. All heatmaps, aggregate and box plots are normalized to a total of 10 million reads.

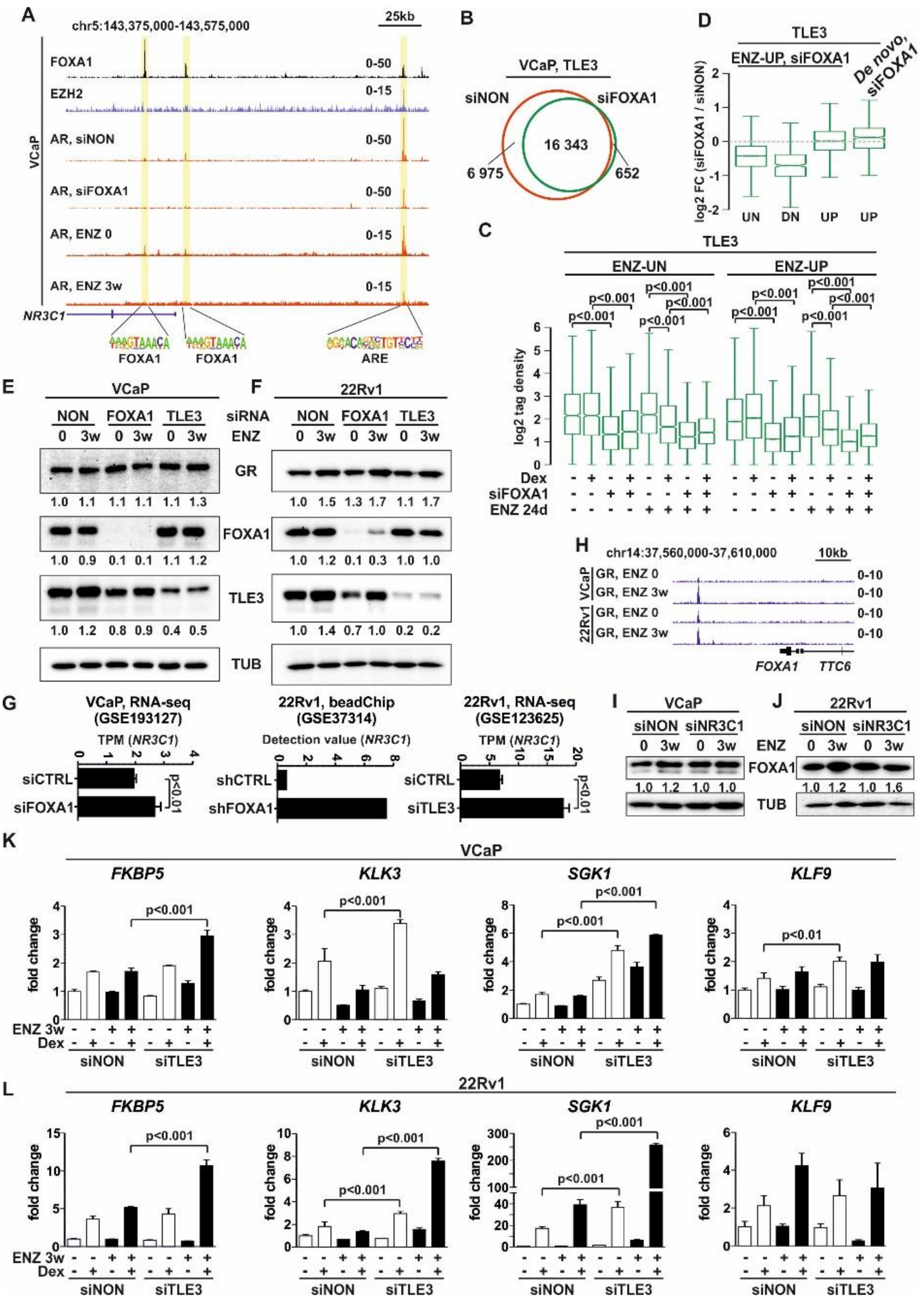


**Supplementary Figure S11. Regulation of NR3C1 by FOXA1.** (A) Genome browser tracks of *NR3C1* loci depicting FOXA1 ChIP-seq, EZH2 ChIP-seq, and AR ChIP-seq in VCaP cells. AR ChIP-seq is shown in cells treated with siNON or siFOXA1, and in ENZ 0 and ENZ 3w cells. Location of FOXA1 and ARE binding motifs are indicated at the bottom of the tracks. (B) Venn diagrams of TLE3 chromatin binding in VCaP cells treated with siNON (red circle) or siFOXA1 (green circle). (C) Box plots represent the normalized log2 tag density of TLE3 ChIP-seq at ENZ-UN and ENZ-UP sites in VCaP cells treated with or without siFOXA1. (D) Log2 fold change (FC) of TLE3 ChIP-seq tags treated with siFOXA1 versus treated with siNON at ENZ-UP siFOXA1-UN, ENZ-UP siFOXA1-DN, ENZ-UP siFOXA1-UP, and *de novo* siFOXA1-UP sites in VCaP cells. (E-F) Immunoblotting of GR, FOXA1, TLE3, and TUB protein levels in VCaP (E) and 22Rv1 (F) ENZ 0 and ENZ 3w cells treated with siNON or siFOXA1 or siTLE3. Quantification of GR, FOXA1, and TLE3 levels to TUB is shown below the immunoblot. (G) Transcript levels of NR3C1 using indicated method in VCaP cells after siFOXA1 (left), in 22Rv1 cells after shFOXA1 (middle), and in 22Rv1 cells after siTLE3 (right). (H) Genome browser tracks of *FOXA1* loci depicting GR ChIP-seq in VCaP and 22Rv1 ENZ 0 and ENZ 3w cells. (I-J) Immunoblotting of FOXA1 and TUB protein levels in (I) VCaP or (J) 22Rv1 ENZ 0 and ENZ 3w cells treated with siNON or siNR3C1. Quantification of FOXA1 levels to TUB is shown below the immunoblot. (K-L) Bar graphs depict GR target gene analysis of *FKBP5*, *KLK3*, *SGK1*, *KLF9* in VCaP (K) and 22Rv1 (L) ENZ 0 and ENZ 3w cells treated with siNON or siTLE3 in the presence or absence of Dex. Bars represent mean ±SD, n=3. All genome browser tracks, and box plots are normalized to a total of 10 million reads. Statistical significance calculated with unpaired t-test (2 conditions) or using One-way ANOVA with Bonferroni *post hoc* test (3 conditions).

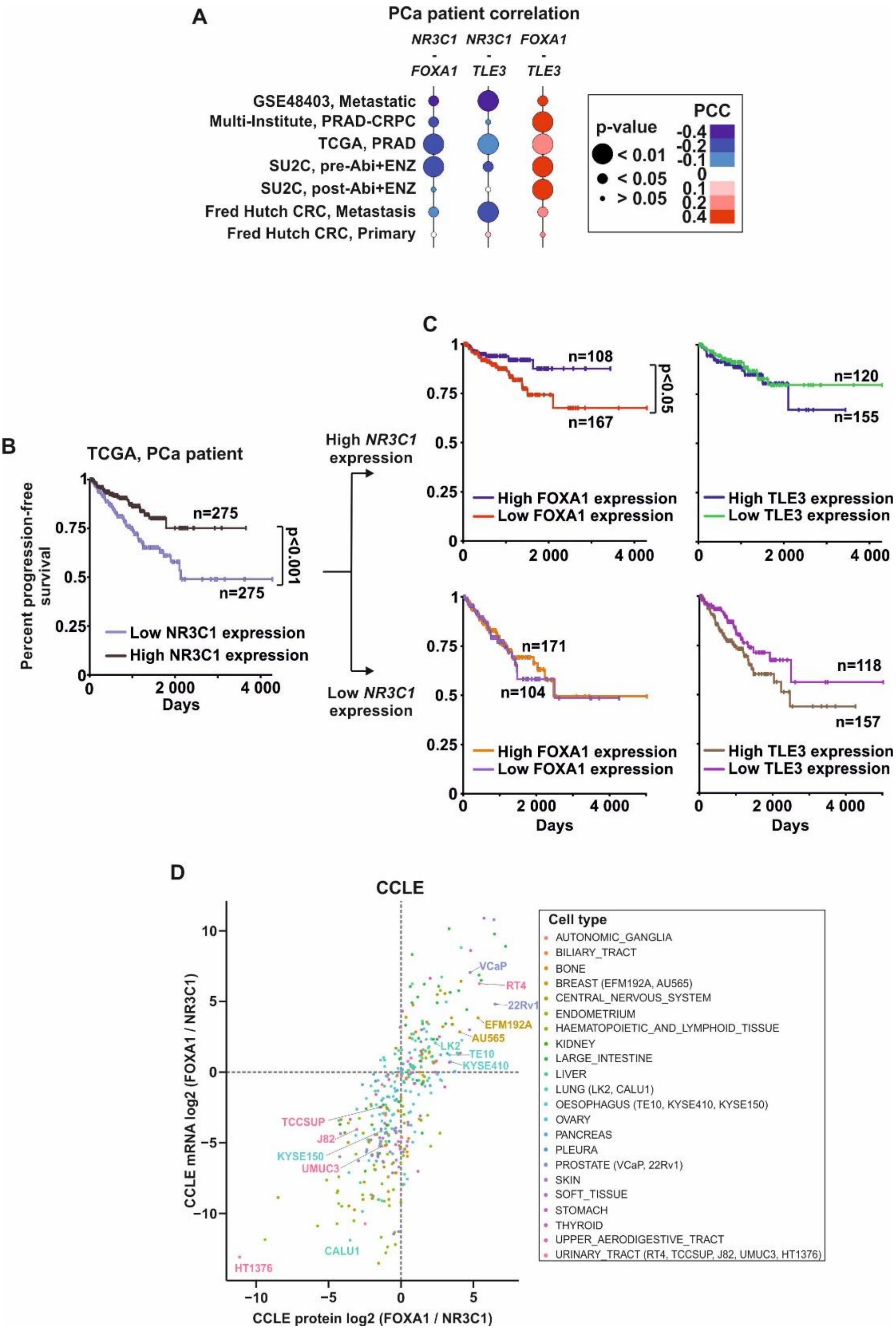


**Supplementary Figure S12. Prostate cancer patient dataset analysis.** (A) Pearson correlation coefficiency (PCC) of *NR3C1*, *FOXA1*, and *TLE3* transcript levels at indicated PCa patient datasets. Circle color represents PCC value with blue indicating negative and red positive correlation. Circle size depicts p-value. (B) Progression-free survival of TCGA PCa patients divided into low (light blue) and high (black) *NR3C1* expression. (C) Progression-free survival of TCGA PCa patients with high (upper) or low (lower) *NR3C1* expression divided into low and high *FOXA1* (left) or low and high *TLE3* (right) expression. Color coding: high *NR3C1* high *FOXA1*, blue; high *NR3C1* low *FOXA1*, red; high *NR3C1* high *TLE3*, dark blue; high *NR3C1* low *TLE3*, green; low *NR3C1* high *FOXA1*, orange; low *NR3C1* low *FOXA1*, purple; low *NR3C1* high *TLE3*, brown; low *NR3C1* low *TLE3*, violet. Statistical significance in the survival graphs is calculated with log-rank test. (D) Log2 fold change of *FOXA1* versus *NR3C1* transcript (y-axis) and protein (x-axis) expression in CCLE cell types. Cell types color coded with anatomical location. Specific cell lines are highlighted in the graph and in the legend.

**Supplementary Table S1. RT-qPCR primers.**

| | |
|---|---|
| *FKBP5* | Forward: 5’-AAAAGGCCAAGGAGCACAAC-3’<br>Reverse: 5’-TTGAGGAGGGGCCGAGTTC-3’ |
| *KLK3* (PSA) | Forward: 5’-GGCAGGTGCTTGTGGCCTCTC-3’<br>Reverse: 5’-CACCCGAGCAGGTGCTTTTGC-3’ |
| *SGK1* | Forward: 5’-ATGCCAACCCTTCTCCTCC-3’<br>Reverse: 5’-TCAACAGAACATTCCGCTCC-3’ |
| *KLF9* | Forward: 5’-TACAGTGGCTGTGGGAAAGT-3’<br>Reverse: 5’-AGCGGGAGAACTTTTTAAGG-3’ |
| *NR3C1* (GR) | Forward: 5’-GAACTGGCAGCGGTTTTATC-3’<br>Reverse: 5’-TGGTATCTGATTGGTGATGATTTC-3’ |
| *RPL13A* | Forward: 5’-ACGACAAGAAAAAGCGGATGG-3’<br>Reverse: 5’-AGGGCAACAATGGAGGAAGG-3’ |

**Supplementary Table S2. RNA-seq data.**

**Supplementary Table S3. Pathway enrichment analyses.**

**Supplementary Table S4. Public datasets utilized in the study.**

**Supplementary Table S5. ChIP-seq peaks.**

**Supplementary Table S6. Motif enrichment analyses.**
